# The Lateral Protein Cluster as a Key Component of Plant Cell Polarity

**DOI:** 10.64898/2026.08.17.745151

**Authors:** Akira Yoshinari, Keito Yunoki, Kei Ota, Karin Futami, Kazuki Motomura, Emi Mishiro-Sato, Reika Isoda, Atsushi Takeda, Jelmer J. Lindeboom, Satoshi Naramoto, Masayoshi Nakamura, Wolf B. Frommer

## Abstract

Cell polarity is an ancient organizing principle across kingdoms. As in animal epithelial cells, plant cells asymmetrically distribute proteins to establish functionally distinct membrane domains. In roots, radial polarity distinguishes inner and outer cell surfaces and supports directional nutrient transport, yet its molecular basis remains poorly understood. Here, we show that the leucine-rich repeat receptor-like kinases CaMRLK and IRK occupy complementary lateral plasma membrane domains in *Arabidopsis thaliana* roots. Polarity-guided proximity labeling identified previously uncharacterized proteins associated with inner-and outer-lateral domains. Clade VII LRR-RLKs, protein *S*-acyltransferases, SICK, IRKI1, and a distinct group of NPH3/RPT2-LIKEs assemble into the Lateral Protein Cluster (LPC) through multivalent interactions. LPC components are conserved across land plants, and disruption of NRL function impairs morphogenesis in Arabidopsis and *Marchantia polymorpha*. Together, these findings establish the LPC as an evolutionarily conserved molecular machinery linking radial cell polarity to plant morphogenesis.

## Introduction

Cell polarity is a universal organizing principle that enables cells to translate molecular asymmetry into directional behavior. Polarized distributions of proteins, lipids, and cytoskeletal structures define where cells grow, divide, secrete, sense external cues, or communicate with neighboring cells^1^. In multicellular organisms, these cellular asymmetries must be coordinated across tissues so that local molecular organization is translated into higher-order architecture^2^. However, the mechanisms that connect cell polarity to tissue-scale architecture remain poorly understood in plants.

In plants, one of the best-characterized forms of cell polarity is apical–basal polarity. PIN-FORMED (PIN) auxin transporters are asymmetrically localized at apical or basal plasma membrane (PM) domains and direct intercellular auxin flow, thereby shaping developmental patterning^3,4^. This polarity is maintained by regulatory proteins, including AGC kinases and NPH3/RPT2-LIKE (NRL) proteins of the NPY clade, which act as molecular scaffolds that restrict lateral diffusion and organize regulatory factors at defined PM regions^5–7^. These studies established local protein assemblies as an important mechanism for stabilizing polar PM domains. Whether comparable higher-order assemblies organize polarity along other cellular axes remains largely unknown.

Lateral polarity, corresponding to the radial body axis and distinguishing inner (proximal) from outer (distal) cell surfaces, represents a distinct and comparatively understudied dimension of plant cell polarity. In Arabidopsis roots, the borate efflux transporter REQUIRES HIGH BORON 1 (BOR1) and the boric acid channel NODULIN 26-LIKE INTRINSIC PROTEIN 5;1 (NIP5;1) occupy inner- and outer-lateral PM domains, respectively^8,9^. The receptor-like cytoplasmic kinase PBL15/SCHENGEN1 (SGN1) localizes to the outer-lateral domain of endodermal cells and is required for Casparian strip formation ^10^. Several leucine-rich repeat receptor-like kinases (LRR-RLKs) also exhibit lateral polarity and regulate root development. KINASE ON THE INSIDE (KOIN) localizes to the inner-lateral PM and is involved in cell division. On the other hand, INFLORESCENCE AND ROOT APICES RECEPTOR KINASE (IRK) displays cell-layer-dependent polarity and shapes the orientation and frequency of specific root cell divisions: IRK localizes to the inner-lateral PM in the epidermis and cortex and to the outer-lateral PM in the endodermis^11,12^. These proteins demonstrate that adjacent lateral cell surfaces are molecularly distinct, yet the molecular machinery that organizes these domains and couples their polarity to cell morphogenesis remains unknown.

Here, we identified an LRR-RLK, CALMODULIN-BINDING RECEPTOR-LIKE KINASE (CaMRLK), as a lateral polarity marker whose orientation is opposite to that of IRK. We exploited this spatial contrast to perform polarity-guided proximity proteomics and identify proteins associated with opposing lateral PM domains. This approach uncovered NPT-clade NRL proteins, protein *S*-acyltransferases (PATs), LRR-RLK VIIs, the cytoplasmic kinase SICK, and IRKI1 as components of a multivalent protein assembly that we term the lateral protein cluster (LPC). LPC components colocalize at lateral PM domains, engage in extensive protein–protein interactions, and organize into higher-order PM microdomains. Disruption of NPT-clade NRLs compromises cell morphogenesis and tissue organization in Arabidopsis and liverwort Marchantia. Together, our findings establish a mechanistic link between lateral PM organization and cell morphogenesis and provide a basis for defining the mechanisms and evolutionary principles underlying plant cell polarity.

## Results

### Polar receptor-like kinases mark distinct lateral plasma membrane domains in root cells

To establish a differential proximity-labeling strategy for lateral PM domains, we sought a membrane protein whose polarity was opposite to that of IRK. Screening LRR-RLKs whose subcellular localization had not been characterized identified CALMODULIN-BINDING RECEPTOR-LIKE KINASE/MATERNAL EFFECT EMBRYO ARREST 62 (CaMRLK/MEE62) as a receptor with the inverse polarity pattern. CaMRLK localized to the outer-lateral PM in the epidermis and cortex, as well as the inner-lateral PM in the endodermis, indicating that CaMRLK polarity switches across cell layers (**Figure 1A**) and is opposite to that of IRK. To quantify CaMRLK polarity, we compared its polarity index with those of an LRR-RLK, BRASSINOSTEROID INSENSITIVE 1 (BRI1) and LOW TEMPERATURE-INDUCED PROTEIN 6B (Lti6b), used as non-polar controls^13^ (**Figures 1B–1D and S1**). The polarity index of CaMRLK was 2.4 ± 0.68 in the epidermis, 1.8 ± 0.36 in the cortex, and 0.6 ± 0.12 in the endodermis. In contrast, BRI1 and Lti6b showed polarity indices close to 1, consistent with largely non-polar PM localization.

**Figure 1.**
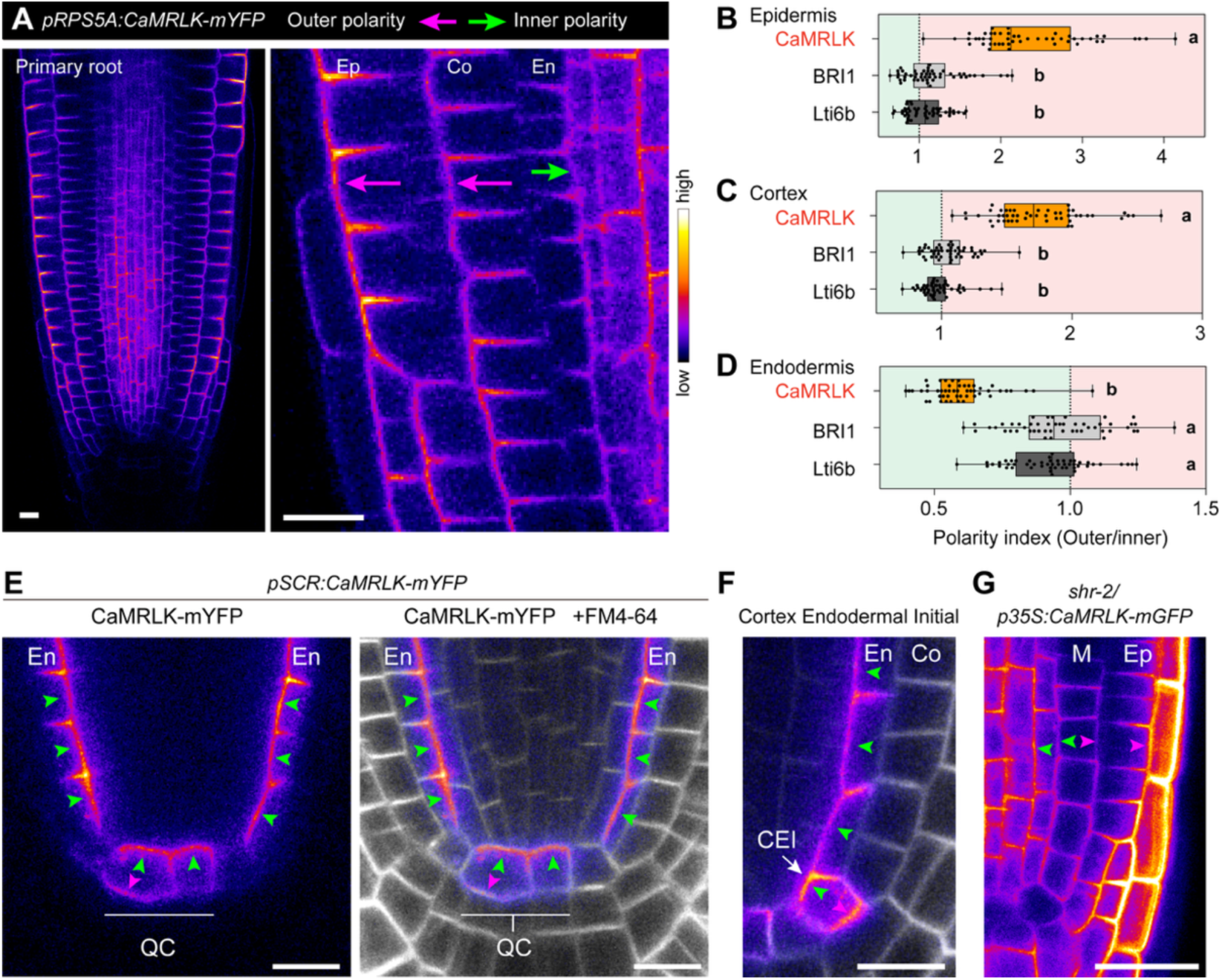
CaMRLK exhibits a dual polarity pattern inverted between endodermis and cortex. (A) Confocal images of 4-day-old *Arabidopsis thaliana* primary root tip harboring *pRPS5A:CaMRLK-mYFP*. Arrows indicate the orientation of CaMRLK-mYFP polarity. Scale bar: 10 µm. (B – D) Polarity index of CaMRLK-mYFP (*pRPS5A:CaMRLK-mYFP*), BRI1-mYFP (*pRPS5A:BRI1-mYFP*), and Lti6b-GFP (*pUBQ10:Lti6b-GFP*) in epidermis (B), cortex (C), and endodermis (D). *n* = 50 cells for each sample. One-way ANOVA with Tukey’s post-hoc test was conducted. Letters denote statistically significant differences, *p* < 0.05. See also **Figure S1**. (E and F) Confocal images of 3-day-old Arabidopsis primary root tip harboring *pSCR:CaMRLK-mYFP* stained with the PM dye FM4-64. Arrows indicate the orientation of CaMRLK-mYFP polarity. Scale bar: 10 µm. (G) Confocal image of a primary root tip of 4-day-old *short-root-2* (*shr-2*) harboring *p35S:CaMRLK-mGFP*. Scale bar: 10 µm. Ep, epidermis; Co, cortex; En, endodermis; QC, quiescent center; CEI, cortex/endodermis initial; M, aberrant cell layer.

To further examine the spatial regulation of CaMRLK polarity, we expressed CaMRLK-mYFP under the SCARECROW (SCR) promoter in the ground-tissue lineage and quiescent center (QC)^14^. CaMRLK-mYFP localized to the inner-lateral PM in differentiated endodermal cells, consistent with its native polarity pattern (**Figures 1E and 1F**). By contrast, CaMRLK displayed bipolar inner- and outer-lateral localization in QC and cortex/endodermal initial (CEI) cells. A bipolar pattern was also observed in the single ground-tissue layer of the *short-root* mutant (*shr-2*)^15^ (**Figure 1G**), indicating that CaMRLK polarity depends on tissue context rather than cell identity alone. Notably, this pattern contrasts with IRK localization in *shr*, where IRK accumulates in the central region of the aberrant ground-tissue layer^11^.

Thus, CaMRLK and IRK define opposing lateral PM domains across root cell layers, providing spatially distinct baits for differential proximity labeling.

### Identification of proteins polarly localized in the lateral plasma membrane domains

Leveraging the opposing polarity of CaMRLK and IRK, we established a spatially resolved proximity-labeling approach to identify proteins associated with distinct lateral PM domains. We generated Arabidopsis lines expressing the engineered biotin ligase TurboID (TbID) fused proteins, CaMRLK-TbID-GFP, IRK-TbID-GFP, or the cytosolic control β-Glucuronidase-GFP-TbID-GFP (GUS-2xGFP), under the control of the *SCR* promoter (**Figure 2A**). In endodermal cells, CaMRLK-TbID-GFP and IRK-TbID-GFP localized to the inner- and outer-lateral PM domains, respectively, while GUS-2xGFP-TbID localized to the cytosol (**Figure 2B**).

**Figure 2.**
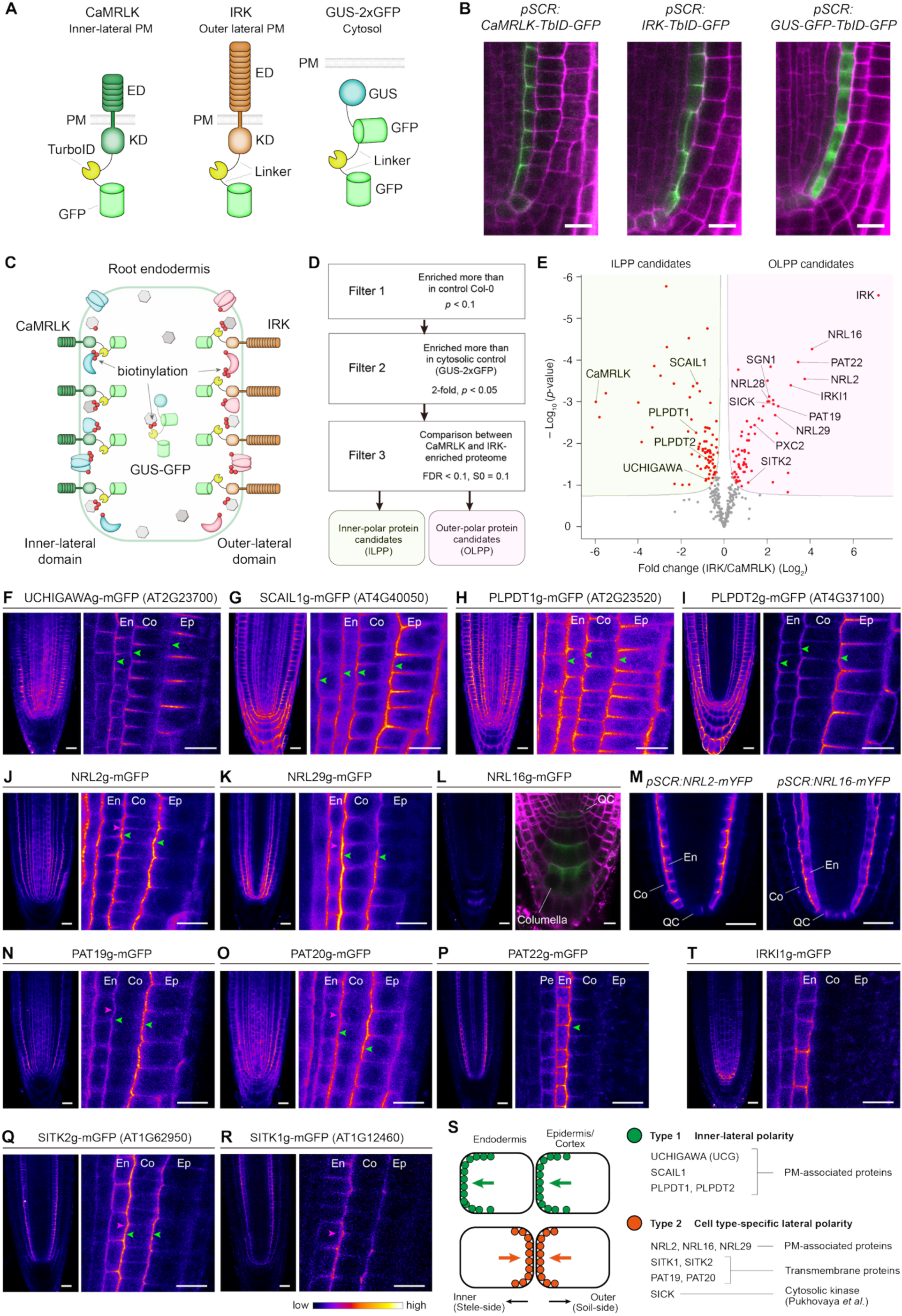
Polarity-guided spatial proteomics identified uncharacterized polar proteins. (A) Schematic representation of the TbID-fused constructs. (B) Confocal images of CaMRLK, IRK, and GUS-GFP fused with TbID-GFP and expressed under the control of the *SCR* promoter. Plants were grown on MGRL medium containing 1 µM B for 9 days. Cell walls were stained with propidium iodide (PI) and are shown in magenta. Scale bar: 10 µm. (C) Conceptual illustration of the polarity-guided spatial proteomics. TbID fused to CaMRLK or IRK preferentially biotinylates proteins localized at the inner or outer lateral PM domains, respectively. By contrast, TbID fused to GUS-GFP is localized in the cytosol and preferentially labels cytosolic proteins. (D) Flowchart of the spatial proteomics workflow. In Filter 1, proteins significantly enriched relative to Col-0 plants lacking TbID expression were selected (*p* < 0.1, determined by Welch’s *t*-test). In Filter 2, proteins enriched more than 2-fold relative to the GUS-GFP control were selected (*P* < 0.05, Welch’s *t*-test). In Filter 3, proteins preferentially enriched in either the CaMRLK-TbID-GFP or IRK-TbID-GFP proteome were identified using Perseus software (FDR < 0.1, S_0_ = 0.1). (E) Volcano plot comparing the CaMRLK-TbID proxitome and the IRK-TbID proxitome. Proteins tested in this study are highlighted in red. Polar proteins are indicated by outlined labels (FDR < 0.1, S_0_ = 0.1). (F – L, N – R) Confocal images of selected ILPPs and OLPPs in the root tips of *Arabidopsis thaliana* transgenic plants grown for 4–5 days. Arrowheads indicate polarity orientation of the proteins. Scale bar: 20 µm (left panels) and 10 µm (right magnified panels). Pe, pericycle; En, endodermis; Co, cortex; Ep, epidermis; QC, quiescent center. Each construct comprised the native promoter and genomic coding sequence, including introns, fused to mGFP and followed by the *NOS* terminator. For L, Primary root was stained with 1 µM FM4-64 to visualize the PM. (M) Confocal images of NRL2-mYFP and NRL16-mYFP expressed under the control of the *SCR* promoter. Scale bar: 20 µm. (S) Summary of the polarity proteins identified in this study and in the accompanying study by Pukhovaya *et al.* (T) Confocal image of IRKI1-mGFP. Scale bar: 20 µm (left panel) and 10 µm (right magnified panel).

We hypothesized that CaMRLK-TbID-GFP and IRK-TbID-GFP would preferentially biotinylate proteins that physically interact with, or reside in close proximity to, the respective fusion proteins, thereby enriching proteins associated with the inner- and outer-lateral PM domains of endodermal cells. Based on their relative enrichment in the CaMRLK- or IRK-proximal proteome (proxitome), we classified these proteins into inner-lateral polar protein (ILPP) candidates and outer-lateral polar protein (OLPP) candidates, respectively (**Figures 2C and 2D**). To reduce nonspecific background, we retained proteins that were significantly enriched relative to both wild-type Col-0 plants lacking TbID constructs and the cytosolic GUS-2xGFP-TbID control. As expected, the CaMRLK and IRK proxitomes were distinct and enriched for PM- and/or cell periphery-associated proteins, whereas the GUS-2xGFP-TbID proxitome was enriched for cytosolic proteins (**Figure S2**). Comparison of the proxitomes identified 159 candidate polar proteins (**Figure 2E; Table S2**). The identification of previously characterized polar proteins, including PIN3^16^ among ILPP candidates and PXC2/CANAR^17^ and SGN1^10^ among OLPP candidates, supported the ability of this approach to resolve lateral PM-associated proteomes. To determine whether other candidates also localize to polar domains, we generated transgenic plants expressing 24 different fluorescent protein fusions and found them to be enriched in the polar proteome across independent analyses.

### Proteins localized in the inner-lateral PM domain

From the 78 ILPP candidates, we evaluated localization of 11 ILPPs, which were identified by independent replicates with high enrichment scores, by confocal microscopy in *Arabidopsis thaliana* roots. AT2G23700 encodes a MIP1 leucine zipper domain and Domain of Unknown Function 547 (DUF547)-containing protein here named UCHIGAWA (UCG; “inner side” in Japanese) (**Figures 2F and S3**); AT4G40050 was named SCAI-LIKE 1 (SCAIL1) because it shares 34% amino acid sequence identity with human SUPPRESSOR OF CANCER CELL INVASION (SCAI) (**Figures 2G and S4**); and two putative pyridoxal 5’-phosphate (PLP)-dependent transferases, designated PLPDT1 (AT2G23520) and PLPDT2 (AT4G37100) (**Figures 2H**, **2I, and S5**), exhibited clear inner-lateral polar PM localization. All four proteins lack obvious transmembrane helices, indicating that they associate with the PM either directly or indirectly via interaction with other PM proteins (**Figures S3–S5**). SCAR1/WAVE1, previously reported to accumulate at cell corners, was also identified and exhibited corner-associated and bipolar localization patterns^18^ (**Figure S6A**).

Not all ILPP candidates were detectably polarized. For example, the PSI-LIKE/DUF668-containing proteins PSI3 and its homologs PSIL-LIKE 1 (PSIL1; AT3G23160), and PSIL3 (AT5G04550), the protein phosphatase 2A (PP2A) B′θ subunit, the MECHANOSENSITIVE CHANNEL OF SMALL CONDUCTANCE-LIKE 4 (MSL4), and a DUF247-containing protein (AT3G47210) fusions localized to the PM but showed no detectable polar distribution (**Figure S6B**), indicating that enrichment in a polar proxitome reflects proximity to the bait-defined membrane environment but does not necessarily imply polarized localization.

### Proteins exhibiting a tissue-specific polar localization in the lateral PM domains

The 65 OLPPs included members of several notable protein families, including NPH3/RPT2-LIKE (NRL), PROTEIN *S*-ACYLTRANSFERASEs (PAT), and LRR-RLK VII including SITK2 and PXC2/CANAR (**Figure 2E**). We examined the subcellular localization of 13 of the 65 OLPPs, prioritizing members of these three protein families as well as candidates, which were identified by independent replicates with high enrichment scores. Among the NRLs identified in the IRK proxitome, NRL2, NRL16, and NRL29 exhibited polarized localization at the lateral PM domains (**Figures 2J–2L**), whereas NRL28 and NPH3 did not (**Figure S6C**). Intriguingly, all three polarized NRLs belong to the same phylogenetic group, previously designated the NPT clade^19^. Furthermore, NRL2 and NRL29 displayed a reversal in polarity orientation across root cell types, similar to IRK (**Figures 2J and 2L**). When expressed under the control of the *SCR* promoter, NRL16 also exhibited outer-lateral polarity in endodermal cells, similar to NRL2 (**Figure 2M**).

PAT19, PAT20, and PAT22, previously characterized as PM-localized Group-C PATs^20^, also exhibited lateral PM polarity (**Figures 2N–2P**). Their polarity orientation similarly reversed between the cortex and endodermis, closely matching that of NPT-clade NRLs and IRK. Polarity of the more weakly expressed PAT22 was independently confirmed using constructs driven by *RPS5A* and *SCR* promoters (**Figure S7**).

A related pattern was observed for LRR-RLK VII proteins. PXC2/CANAR, the closest homolog of IRK, had previously been reported to localize laterally^17^, and SITK2 (AT1G62950) likewise exhibited cell-layer-dependent lateral polarity (**Figures 2Q**). Although its closest homolog AT1G12460 (SITK1), was not detected in the IRK proxitome, SITK1 exhibited lateral PM polarity similar to that of SITK2 (**Figure 2R**). Together, the NPT-clade NRLs, PATs, and SITK1/2 localized to the inner-lateral PM domain in the epidermis and cortex and to the outer-lateral PM domain in the endodermis, as well as IRK and PXC2 (**Figures 2S**).

As for ILPPs, OLPP enrichment could not be confirmed for all candidates, e.g. IRK-INTERACTING 1 (IRKI1), previously referred to as IRKI ^21^ did not exhibit detectable polarity when expressed from either its native or the *RPS5A* promoter (**Figures 2T and S6C**). Casein kinase 1-like 7 (CKL7) showed a polarized distribution; however, it preferentially localized to the cytosol and apical–basal PM domains rather than to the lateral PM domains, and thus did not recapitulate the outer-lateral polarity expected for an OLPP (**Figure S6C**). The receptor-like cytoplasmic kinases RLCK-VI B1 and PBL35 localized to the PM but showed no detectable polar distribution (**Figure S6C**). We were unable to detect fluorescence from mGFP-tagged SICK (AT2G40980), a putative cytosolic kinase, in our system. However, Pukhovaya et al. independently showed that SICK exhibits polar localization at lateral PM domains (*Pukhovaya et al., accompanying manuscript)*. We therefore concluded that SICK represents a *bona fide* polarity protein found in the IRK proxitome (**Figure 2S**).

### NPT-clade NRLs localize to the lateral plasma membrane domains

The lateral polarity of NRL2, NRL16, and NRL29 was particularly intriguing because NPY-clade proteins, a subgroup of NRLs, function as molecular scaffolds for PIN auxin transporters and maintain polar localization at apical and basal PM domains^5–7^. We therefore asked whether NPT-clade NRLs constitute a distinct group of NRLs associated with lateral polarity. The NPT clade comprises six members, NRL2, NRL16, NRL19, NRL22, NRL24, and NRL29 (**Figure 3A**). In roots, NRL2 showed broad protein accumulation, whereas NRL16, NRL19, and NRL29 exhibited more cell-type-specific accumulation (**Figures 2J–2L**, **3B–3E**). By contrast, NRL22 and NRL24 accumulated predominantly in reproductive organs rather than in roots, consistent with their transcript accumulation patterns (**Figures 3F–3M and S8**). Notably, NRL19 was not detectably accumulated in root endodermal cells, the cell type in which TbID-based proximity labeling was performed (**Figure 3N**), consistent with its absence from the resulting proxitome. Together, these results indicate that NPT-clade NRLs share a characteristic association with lateral polarity.

**Figure 3.**
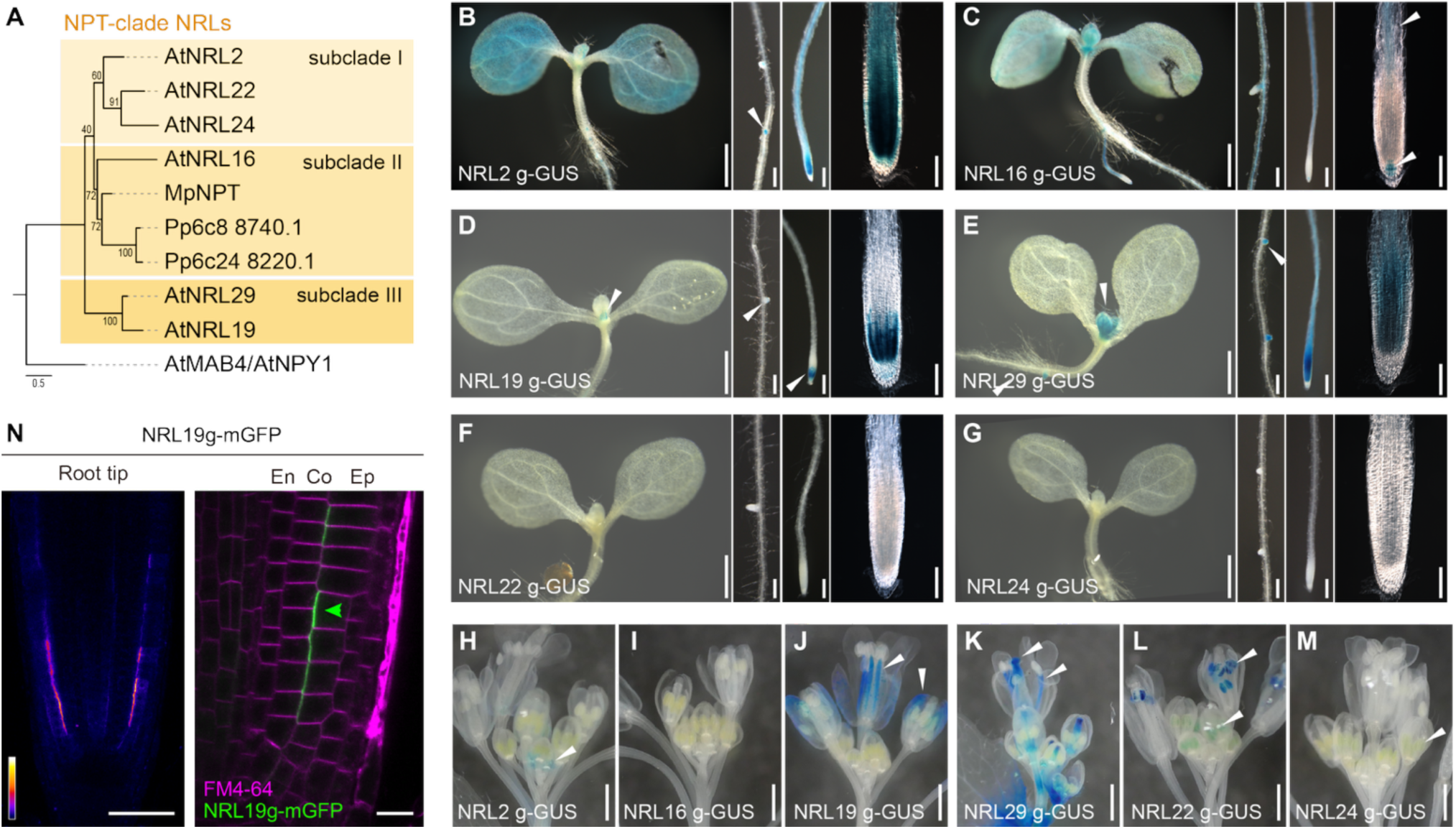
NPT-clade NRLs are polarized in the lateral PM domains. (A) Maximum-likelihood phylogenetic tree of NPT-clade NRLs from *Arabidopsis* (NRL2, NRL16, NRL19, NRL22, NRL24, and NRL29), together with MpNPT from *Marchantia polymorpha* and close homologs from *Physcomitrium patens*. *A. thaliana* MAB4/NPY1/NRL20 was used as the outgroup. NPT proteins were classified into three subclades I–III. Bootstrap values (%) are indicated at the nodes. (B – M) Histochemical analysis of translational GUS reporter lines for NPT-clade NRLs in Arabidopsis. Each construct contained the native promoter and genomic coding sequence, including introns, fused to GUS. (B – G) Microscopy of GUS reporters in seedlings grown for 6 days on MGRL medium. Arrows indicate GUS reporter-positive tissues. Scale bar: 100 µm (left panels; stereomicroscopy), 200 µm (middle two panels; stereomicroscopy), and 100 µm (right panels; transmitted light microscopy). (H – M) Photographs of GUS reporters in floral organs from plants grown in soil for 30 days. Scale bar: 10 mm. (N) Confocal image showing the localization of NRL19-mGFP expressed under the control of its native promoter. The arrow indicates the direction of polarity. Scale bars: 50 µm (left) and 10 µm (right, magnified view).

The *Arabidopsis* genome encodes 33 NRL homologs, which can be broadly classified into seven phylogenetic groups: the NPY (8 members), NPH3 (4 members), NCH1 (8 members), NPS (3 members), NPE (2 members), NPT (6 members), NRL9/NRL25 (2 members) clades^19^. NRLs share a common domain architecture, comprising an N-terminal Broad-complex, Tramtrack, Bric-a-brac (BTB)/poxvirus and zinc finger (POZ) domain, a central plant-specific NPH3 domain, and α-helical and unstructured regions in the C-terminal region (**Figure S9**). Considering the role of NPY-clade NRLs in maintaining apical–basal polarity, two key questions arise: what are the functions of NPT-clade NRLs at the lateral PM domains, and do they act as molecular scaffolds similar to NPY-clade NRLs?

### Identification of Lateral Protein Cluster (LPC)

If the NPT-clade NRLs function as scaffolds of a protein cluster, they physically interact with other components of the protein cluster. To identify the proteins associated with NRL2, an abundantly accumulated NPT member in the seedling (**Figure 3B**), we performed co-immunoprecipitation (co-IP) of NRL2-mYFP and YFP-Lti6b, a non-polar PM-resident protein, from root extracts, followed by comparative LC-MS/MS analysis (**Figure 4A**). Strikingly, several OLPPs were also identified with high confidence in the NRL2 interactome, including three other NPT-clade NRLs (NRL16, NRL19, and NRL29), the polar PATs PAT19, PAT20, and PAT22; the LRR-RLK VIIs SITK1/2, IRK, and PXC2; as well as SICK and IRKI1 (**Figure 4A**). To further validate these associations, we performed reciprocal co-IP using PAT19-mGFP and IRKI1-mGFP as baits and confirmed that most of those proteins were co-immunoprecipitated with both PAT19 and IRKI1 (**Figures 4B–4E**). Together, these results indicate that NRL2, PAT19, IRKI1, and LRR-RLK VII proteins form a *bona fide* protein cluster in close proximity to IRK (**Figure 4E**).

**Figure 4.**
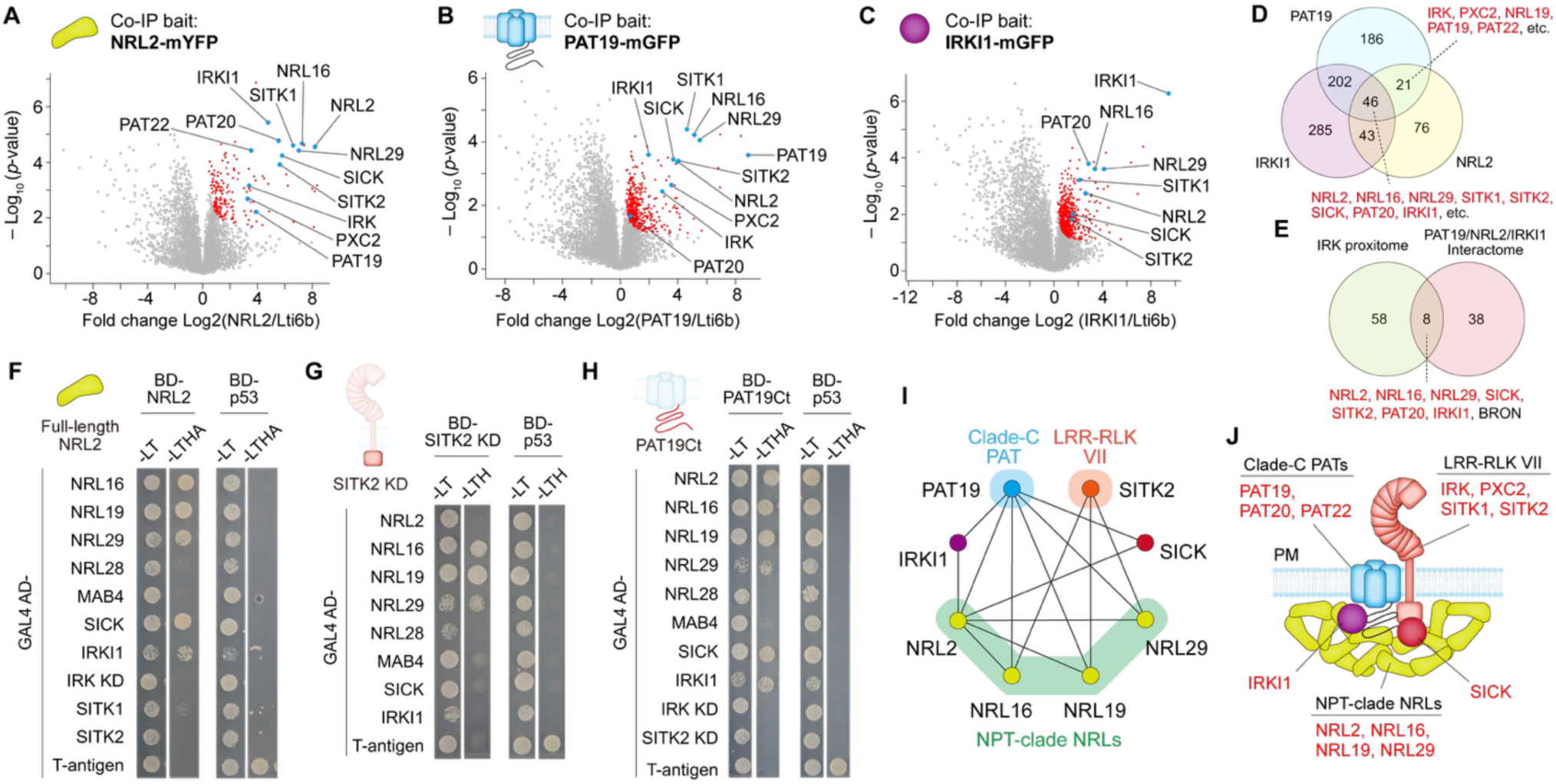
Identification of lateral protein cluster (LPC). (A – C) Volcano plots of enriched proteins in co-immunoprecipitation (co-IP) using YFP-Lti6b (control PM protein) and either NRL2-mYFP, PAT19-mGFP, or IRKI1-mGFP (FDR < 0.1, S_0_ = 0.1). Proteins enriched in the TbID-based proxitome were highlighted. (D) Venn diagram of proteomes enriched as interacting protein candidates with either NRL2-mYFP, PAT19-mGFP, or IRKI1-mGFP (FDR < 0.1, S_0_ =0.1). (E) Venn diagram of proteomes similarly enriched by co-IP and TbID-based IRK proxitome. (F – I) Yeast two-hybrid (Y2H) assays between the potential LPC components. p53 and SV40 large T-antigen were used as the bait and prey, respectively, for the positive control pair. -LT, minimal medium lacking L-Leucine and L-Tryptophan; -LTH, minimal medium lacking L-Leucine, L-Tryptophan, and L-Histidine; -LTHA, minimal medium lacking L-Leucine, L-Tryptophan, L-Histidine, and adenine. (F) Y2H assay between NRLs. Full-length proteins were used. (G) C-terminal kinase domain (KD) (Leu-538 to Ser-890, immediately downstream of the predicted transmembrane domain) of SITK2 was used as the bait for Y2H assay. (H) C-terminal tail fragment (Pro-379 to Lys-718) of PAT19 was used as the bait for Y2H assay. (I) Summary of Y2H assay performed in this study. (J) Schematic illustration of the hypothetical structure of the lateral protein cluster (LPC).

To validate physical interactions among components of the putative protein cluster, we performed yeast two-hybrid (Y2H) assays. We examined the interaction of NRL2 with representative members of the NPT clade (NRL16, NRL19, and NRL29), the NCH1 clade (NRL28, detected in the IRK proxitome), and the NPY clade (MAB4) (**Figures 4F and S10A**). NRL2 interacted strongly with NRL16, NRL19, and NRL29, whereas its interaction was weaker with NRL28 and MAB4 (**Figure 4F and S10A**). NRL2 also interacted with SICK and IRKI1 but not with the cytoplasmic kinase domains of IRK or SITK2 (**Figures 4F and S10A**). We therefore investigated whether these kinase domains interact with other NPT-clade NRLs. Using the SITK2 kinase domain as bait, we found that SITK2 interacted with NRL16, NRL19, and NRL29, but not with NRL2 (**Figures 4G and S10B**). In addition, IRKI1 was previously shown to interact with the kinase domain of IRK^21^. These findings indicate that different components establish distinct interactions within the protein cluster and that IRKI1 may connect to the cluster through the cytoplasmic domains of IRK and potentially those of the related LRR-RLK VIIs PXC2/CANAR and SITK1/2.

In Arabidopsis, PAT19, PAT20, PAT21, and PAT22 had been reported to belong to the same clade and be PM localized ^20,22,23^. These PATs are distinguished by relatively long C-terminal regions containing extensive intrinsically disordered regions (IDR; **Figure S11A**)^20^. To determine whether the C-terminus contributes to polar localization, we analyzed PAT19 variants carrying progressive C-terminal truncations (Δ651–718, Δ609–718, and Δ389–718; **Figures S11B and S11C**). Truncation of the C-terminal region resulted in a reduction in polarity (**Figure S11C**), demonstrating that the PAT19 C-terminal region is important for its polar localization. We therefore hypothesized that the C-terminal region promotes PAT19 polarity by serving as a protein interaction platform within the cluster. Consistent with this hypothesis, Y2H assays showed that the PAT19 C-terminal region (residues 379–718, 340 aa) directly interacted with NRL2, NRL16, NRL19, NRL29, SICK, and IRKI1, but not with NRL28, MAB4, or the cytoplasmic domains of IRK or SITK2 (**Figure 4H and S10C**). Together, our results demonstrate that NRL2 and PAT19 engage in multivalent protein–protein interactions (PPI; **Figure 4I**), which may facilitate the assembly of a multicomponent protein cluster at lateral PM domains. We named this cluster, comprising NPT-clade NRLs, PATs, LRR-RLK VIIs, SICK, and IRKI1, the lateral protein cluster (LPC; **Figure 4J**).

### LPC components are colocalized at the lateral PM domains

To determine whether LPC components colocalize *in vivo*, we performed confocal microscopy. In Arabidopsis root meristems, at the inner-lateral surface of epidermal cells, NRL2-mRFP, NRL29-mGFP, PAT19-mGFP, and SICK-tdTomato exhibited punctate patterns and colocalized with each other (**Figures 5A–5G**). Because IRKI1 and the LRR-RLK VIIs were expressed at low levels at the inner-lateral surface of epidermal cells, we employed a heterologous overexpression system using tobacco *N. benthamiana* leaf epidermal cells (**Figure 5H**). In this system, IRKI1-mCherry, IRK-mGFP, and NRL2-StayGold-E138D (mSG) or -mCherry formed puncta even at the outer surface of leaf epidermal cells, despite this domain being geometrically opposite to the inner-lateral surface where these proteins preferentially localize in Arabidopsis root epidermal cells (**Figures 5I and 5J**). The puncta of IRK-mGFP partially overlapped with those of NRL2-mCherry and IRKI1-mCherry showed a clear colocalization with NRL2-mSG (**Figures 5I–5L**). Together, these results demonstrate that LPC components colocalize in punctate PM structures *in vivo* and establish the heterologous expression system in tobacco as a useful platform for resolving their subcellular organization at high spatial resolution.

**Figure 5.**
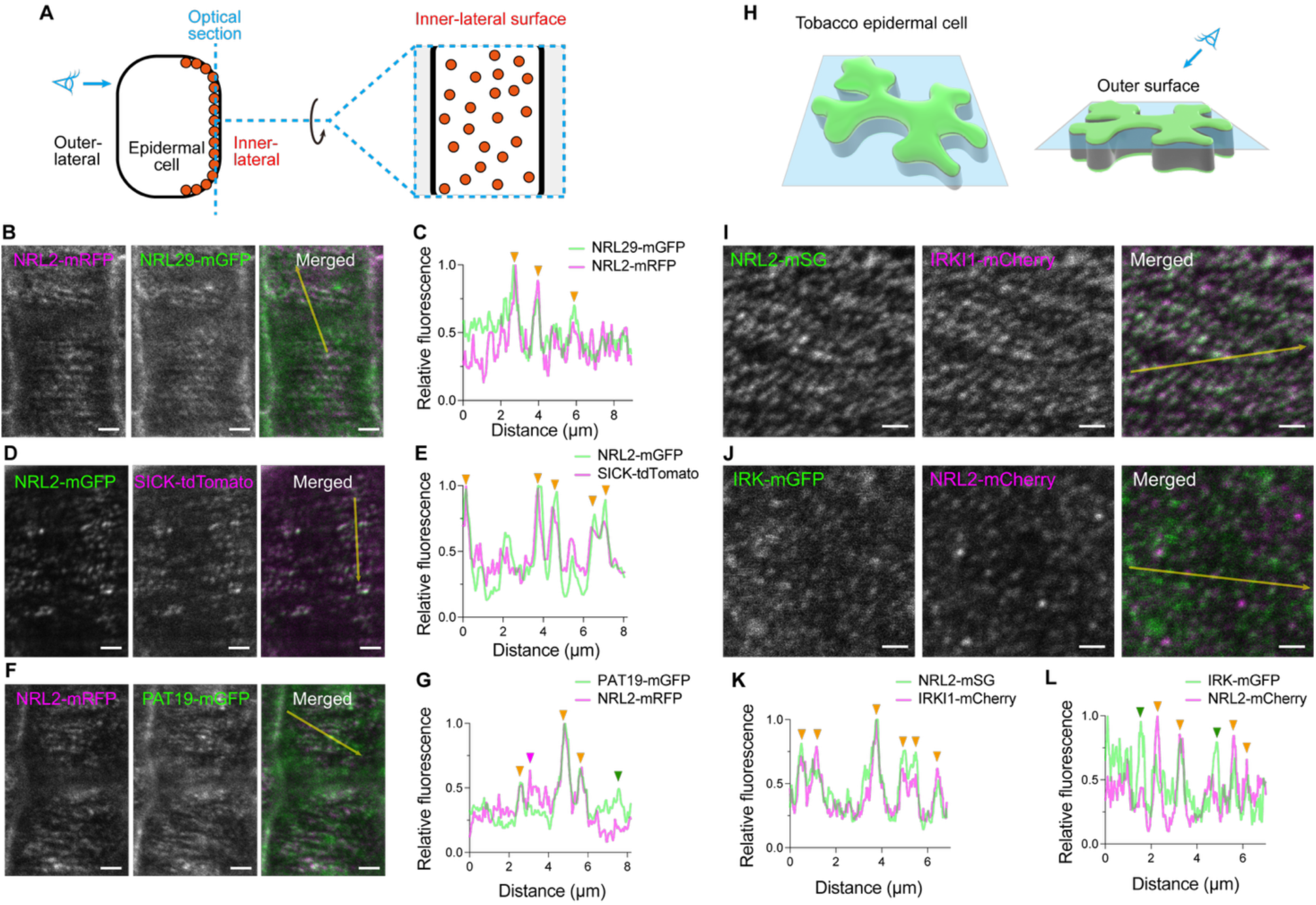
LPC components colocalize in vivo. (A) Schematic illustration of a root epidermal cell. Inner-lateral surface of epidermal cells was observed by confocal microscopy in B, D, and F. (B, D, and F) Confocal images of the inner-lateral surface of Arabidopsis root epidermal cells co-expressing NRL2-mRFP and NRL29-mGFP (B), NRL2-mGFP and SICK-tdTomato (D), and NRL2-mRFP and PAT19-mGFP (F). Four-day-old seedlings of T_2_ generation were observed. Scale bar: 2 µm. (C, E, and G) Fluorescence intensity along the arrows with 3-pixel width indicated in the confocal images (B, D, and F). Orange arrowheads indicate colocalized foci. Magenta and green arrowheads indicate independent foci. (H) Schematic illustration of a tobacco leaf epidermal cell. The outer surface of the epidermal cells was observed by confocal microscopy in I and J. (I and J) Confocal images of the outer surface of tobacco epidermal cells co-expressing NRL2-mSG and IRKI1-mCherry (I) and IRK-mGFP and NRL2-mCherry (J). Scale bar: 1 µm. (K and L) Fluorescence intensity along the arrows with 3-pixel width indicated in the confocal images in I and J. Orange arrowheads indicate colocalized foci. Magenta and green arrowheads indicate independent foci.

### PAT-dependent *S*-acylation is required for LPC microdomain formation

To further characterize the LPC organization, we analyzed the colocalization of PAT19-mGFP and NRL2-mCherry in the tobacco expression system. When expressed individually, both proteins exhibited punctate localization at the outer PM (**Figures 6A and 6B**). Strikingly, their co-expression induced an interconnected, labyrinthine PM pattern in which PAT19-mGFP and NRL2-mCherry extensively colocalized. By contrast, co-expression of PAT19-mGFP with free mCherry did not induce this pattern (**Figures 6C and 6D**). Closer examination revealed that regions of high protein accumulation formed broad band-like structures (**Figure 6E**). We hereafter refer to these higher-order protein assemblies at the PM as LPC microdomains. Notably, the labyrinthine pattern contained narrow, filament-like gaps from which LPC microdomains were excluded (**Figure 6F**). The morphology of these gaps indicated a possible relationship with cortical microtubules (CMTs), a prominent cytoskeletal component of plant cells. Indeed, CMTs visualized using mScarlet-TUB6 were largely excluded from regions occupied by LPC microdomains (**Figures 6F and 6G**). Thus, LPC microdomains and CMTs exhibit a striking complementary spatial organization at the PM, raising the possibility of a functional relationship between these two systems.

**Figure 6.**
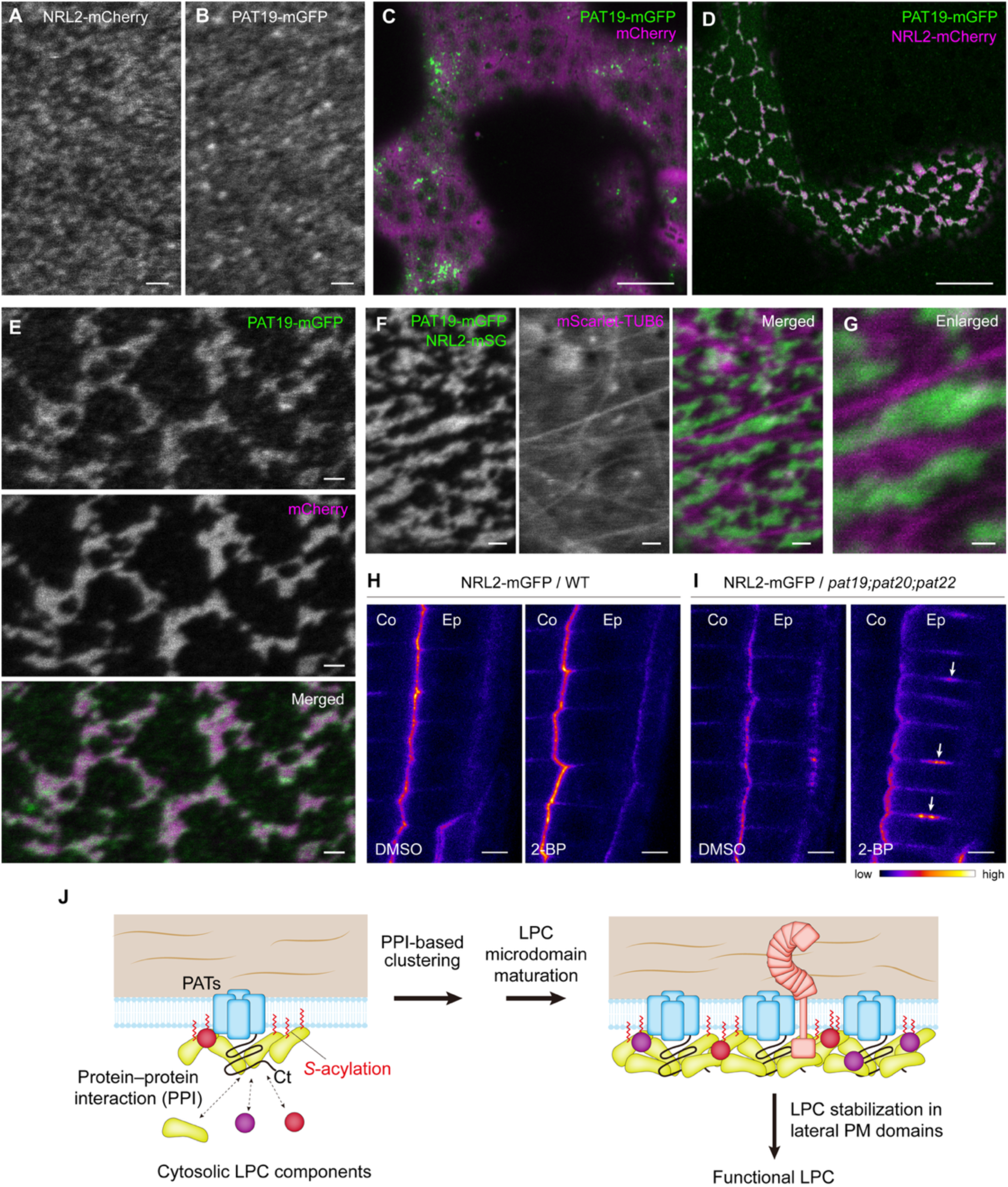
S-acylation is key to LPC microdomain formation. (A and B) Epidermal cells expressing either NRL2-mCherry or PAT19-mGFP under the control of the *UBQ10* promoter. Scale bar: 1 µm. (C and D) Epidermal cells co-expressing PAT19-mGFP and either free mCherry or NRL2-mCherry. Scale bar: 10 µm. (E) High magnification images of epidermal cells co-expressing PAT19-mGFP and NRL2-mCherry. Scale bar: 1 µm. (F and G) Epidermal cells co-expressing PAT19-mGFP, NRL2-mSG, and mScarlet-TUB6. Scale bar: 1 µm (F) and 0.5 µm (G). (H and I) Epidermal cells of *A. thaliana* primary root tips expressing NRL2-mGFP in wild-type or *pat19;pat20;pat22* treated with DMSO (control) or 20 µM 2-bromopalmitate (2-BP) for 14 h. Arrows indicate mislocalization of NRL2-mGFP. Scale bar: 5 µm. (J) Schematic illustration of a hypothetical model of *S-*acylation-dependent microdomain formation. NRLs are likely able to associate with the PM without PATs. *S-*acylation of NRLs and LPC components by PATs enhances the stability of PAT–NRL clustering at PM.

What drives the formation of LPC microdomains? To define the basis of LPC microdomain assembly, we examined the PAT19 C-terminal tail, which binds NRL2 and is required for PAT19 polarity (**Figures 4H and S11**). PAT19^Δ389–718–mGFP^, whose C-terminal tail is deleted, failed to colocalize with NRL2 and abolished LPC microdomain formation in tobacco epidermal cells (**Figure S12**), demonstrating the essentiality of the C-terminal region for microdomain formation. Because PATs are DHHC-type protein *S*-acyltransferases, we hypothesized that PAT-mediated *S*-acylation reinforces the PM association of NRL2 and thereby promotes the formation of LPC microdomains.

To test whether *S*-acylation is required for polar localization of NRL2, Arabidopsis seedlings were treated with 2-bromopalmitate (2-BP), an inhibitor of protein *S*-acylation, for 14 h. In wild-type plants, orientation of the NRL2-mGFP polarity was not altered by 2-BP treatment (**Figure 6H**). Consistently, NRL2-mGFP showed polar localization in *pat19;pat20;pat22* mutants^23^ (**Figure 6I**). Together, these results indicate that PAT19/PAT20/PAT22-dependent *S*-acylation is dispensable for the PM association and polar localization of NRL2 under normal conditions. Strikingly, however, treatment of the *pat19;pat20;pat22* triple mutant with 2-BP caused NRL2-mGFP to mislocalize to the central region of the horizontal PM plane in epidermal cells (**Figure 6I**). These findings indicate two non-mutually exclusive possibilities: *S*-acylation of NRL2 is required for its accurate targeting within the PM, and broad depletion of protein *S*-acylation disrupts the cellular geometric cues that define polar PM domains.

Together, these results support a model in which NRLs associate with the PM independently of *S*-acylation, whereas PAT-dependent *S*-acylation promotes the proper targeting of NRL2 to polar PM domains and drives LPC microdomain assembly and maturation. Reciprocal protein–protein interactions between PATs and NRLs may reinforce this process through a positive-feedback mechanism that stabilizes LPC microdomains (**Figure 6J**).

### Evolutionarily conserved LPC module underlies tissue organization in land plants

The observation that the LPC localizes to lateral PM domains and can form microdomains indicates that its activity is spatially regulated, and may even provide positional information that defines specific PM domains. Consistent with the functional importance of LPC-associated proteins, previous studies reported that mutations in PAT19/PAT20/PAT22, IRK, or PXC2 cause defects in cell morphology in the root meristem^11,17,23^. We therefore asked whether NPT-clade NRLs, which are thought to function as scaffolds within the LPC through multivalent protein–protein interactions, are also required for plant morphogenesis.

To investigate their physiological functions, we analyzed T-DNA insertion mutants of the NPT-clade *NRL* genes (**Figure S13A**). Although single mutants did not show dramatic developmental defects (**Figure S13B**), the *nrl2-2;nrl16-2* double (dKO) and *nrl2-2;nrl16-2;nrl29-1* triple (tKO) mutants exhibited an increased incidence of morphological abnormalities in the root apical meristem (**Figures 7A–7C**). In particular, cellular organization surrounding the quiescent center (QC) was severely disrupted (**Figure 7B**). We also attempted to generate a sextuple NPT-clade NRL mutant by genome editing. However, we were unable to generate null mutants of all the *NPTs*. We therefore generated a line combining the *nrl2;nrl16;nrl19;nrl24;nrl29* quintuple mutant (qKO) with an NRL22 allele carrying an in-frame nine-amino-acid deletion. This line displayed a markedly exacerbated root phenotype, characterized by severe disorganization of the root meristem and abnormal root elongation (**Figures 7A–7E**). Together, these results demonstrate that NPT-clade NRLs act complementarily to maintain root meristem organization and coordinate root cell morphogenesis.

**Figure 7.**
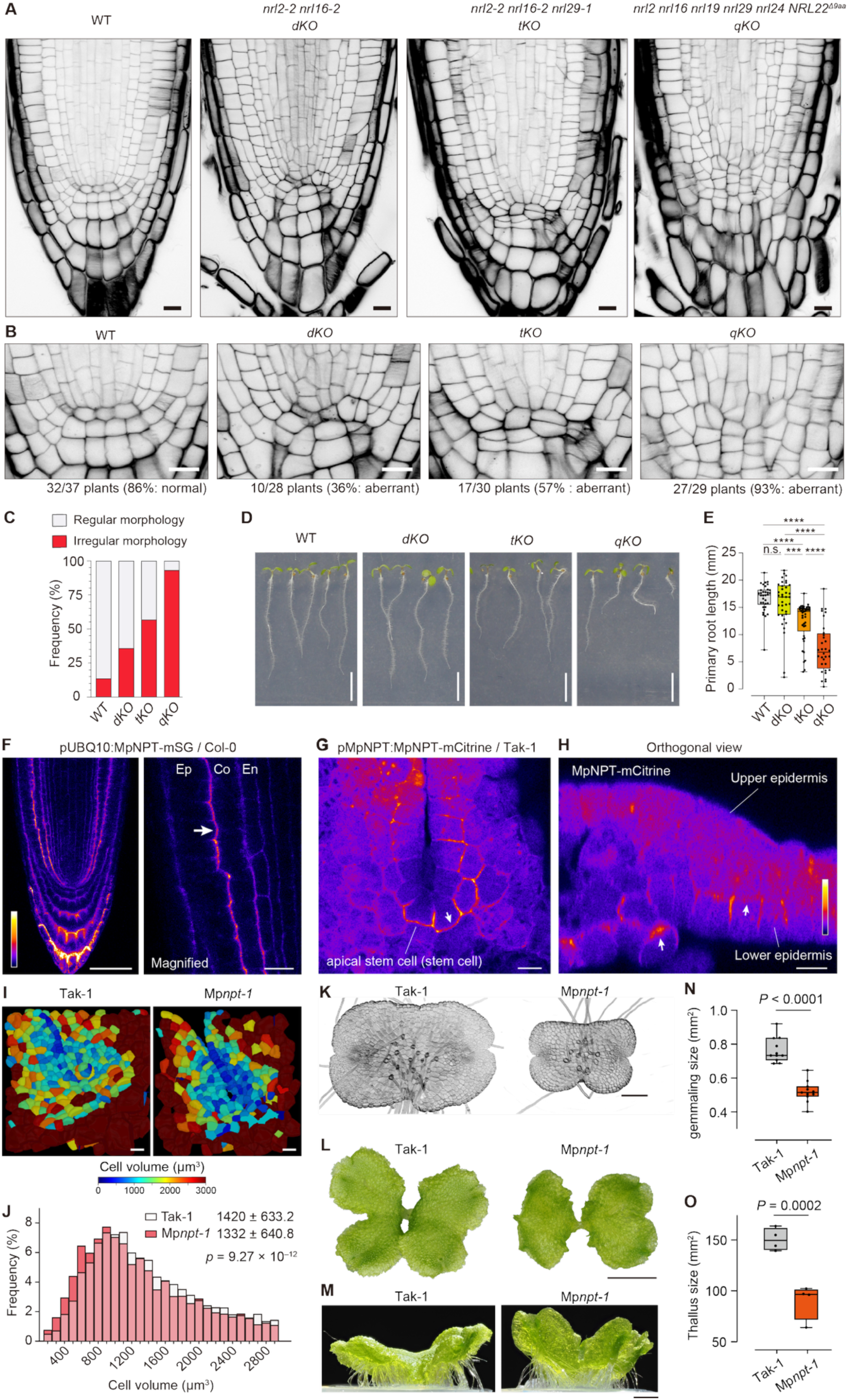
Morphogenetic role and evolutionary significance of NPT-clade NRLs. (A) Confocal images of primary root tips of 3-day-old wild-type and mutant seedlings of *A. thaliana*. Plants were cleared by ClearSee and cell walls were stained with 0.1% (v/v) SR2200 fluorescent dye. Scale bar: 10 µm. (B) Magnified images of stem cell regions. Scale bar: 10 µm. (C) Frequency (%) of roots with regular and irregular morphogenesis in the root apical meristems. *n* = 37 (WT), 28 (dKO), 30 (tKO), and 29 (qKO). (D) Five-day-old seedlings grown vertically on MGRL medium. Scale bar: 5 mm. (E) Box plots for primary root length. Box plot center lines show the medians. Box limits indicate the 25th and 75th percentiles. Bars extend to the minima and maxima. Statistical significance was determined by Tukey’s multiple comparison; n.s. represents *p* > 0.01, \*\*\**p <* 0.001*, ****p <* 0.0001. (F) Confocal image of MpNPT-mSG expressed in *A. thaliana* Col-0 under the control of the *UBQ10* promoter. Scale bar: 50 µm (left) and 10 µm (right, magnified image). Ep, epidermis; Co, cortex; En, endodermis. (G and H) Confocal images of 2-day-old *M. polymorpha* gemmalings expressing MpNPT-mCitrine under the control of the native promoter. Apical meristem notch (G) and an orthogonal view reconstructed from a z-stack (H). Scale bar: 10 µm. (I) Morphological analysis of apical notch cells of WT Tak-1 and Mp*npt-1* using MorphoGraphX software. Cells were colorized based on the volume. Scale bar: 20 µm. (J) Histogram of cell volumes in the apical notch region of wild-type Tak-1 and Mp*npt-1*, plotted using 100 µm^3^ bins. Cell volumes ≤ 200 µm^3^ and ≥ 3,000 µm^3^ were excluded from statistical analysis. *n* = 5797 (Tak-1) and 4198 (Mp*npt-1*) cells from 11 plants each. The *p*-value was determined by two-tailed Welch’s *t*-test. (K) Confocal microscopy of 2-day-old gemmalings of *M. polymorpha* wild-type Tak-1 and Mp*npt-1* grown on Gamborg B5 medium. Cell walls of cleared specimens were stained with SR2200 fluorescent dye. Scale bar: 200 µm. (L) Top-view photographs of 16-day-old thalli of *M. polymorpha* wild-type Tak-1 and Mp*npt-1* grown on Gamborg B5 medium. Scale bar: 2 mm. (M) Side-view photographs of 16-day-old thalli of *M. polymorpha* wild-type Tak-1 and Mp*npt-1* grown on Gamborg B5 medium. Scale bar: 2 mm. (N) Quantification of size of the 2-day-old gemmalings. *n* = 11. *p-*values were determined by two-tailed Welch’s *t*-test. (O) Quantification of size of the 16-day-old thalli. *n* = 4. *p-*values were determined by two-tailed Welch’s *t*-test.

The NPT clade is represented by a single gene in the liverwort *Marchantia polymorpha*, previously designated MpNPT ^19^. To determine whether MpNPT retains the capacity to localize to lateral PM domains, we first examined MpNPT-mSG expressed in Arabidopsis. MpNPT-mSG was clearly enriched at lateral PM domains (**Figure 7F**), indicating that the ability of NPT proteins to target lateral PM regions is evolutionarily conserved across land plants. MpNPT-mCitrine expressed under the control of its native promoter predominantly accumulated in meristematic cells at the apical notch and lower epidermis (**Figure 7G**), whereas its transcript was detected more broadly across the thallus (**Figure S15**). Intriguingly, MpNPT-mCitrine was preferentially enriched at the inner PM domain in the meristematic cells (**Figures 7G and 7H**). This result indicates that the radial polarity axis observed in vascular plants corresponds to the inner–outer polarity of Marchantia, indicating that this polarity framework was already established in an early-diverging land plant lineage. Together, these observations indicate that MpNPT possesses an evolutionarily conserved intrinsic capacity for polar PM localization.

To examine the function of NPT in Marchantia, we generated Mp*npt* mutants and analyzed their developmental phenotypes (**Figures 7I–7O and S16**). Analysis of the apical notch region revealed reduced cell volume in the meristematic zone of Mp*npt-1*, indicating a defect in cell division or proliferation (**Figures 7I and 7J**). The mutants also showed reduced size as gemmalings and mature thalli, together with pronounced upward curvature of the thalli (**Figures 7K–7O**). These cellular and macroscopic defects indicate that MpNPT is required for normal thallus morphogenesis. Together with the developmental defects observed in Arabidopsis NPT-clade mutants, these findings support an evolutionarily conserved role for NPT-clade NRLs in cellular morphogenesis and tissue organization across land plants.

## Discussion

Throughout evolution, cell polarity has enabled living organisms to convert molecular asymmetry into multicellularity and, ultimately, complex tissues and organs ^24^. Because multicellularity evolved independently in the plant and animal lineages, plants likely developed distinct mechanisms for establishing and interpreting cell polarity^25^. Studies of plant cell polarity have yielded detailed insights into apical–basal polarity, particularly through analyses of PIN-mediated auxin transport, and into tip growth in root hairs and pollen tubes^26,27^. By contrast, the mechanisms and molecular mediators that control polarity along the inner–outer radial axis of tissues remain poorly understood^28,29^.

In this study, we established a polarity-guided proximity proteomics platform and generated the first broad molecular inventory of proteins associated with distinct lateral PM domains along the radial body axis (**Figure 2**). This represents a major advance because only a small number of laterally polarized proteins have previously been characterized in plant cells. Among the inner-lateral PM proteins identified in the CaMRLK proxitome, UCG and SCAIL1 are particularly notable because they lack transmembrane domains and canonical lipid-modification motifs and therefore could not have been reliably predicted as PM-associated proteins by conventional sequence-based approaches. Our results demonstrate that proximity labeling can uncover an otherwise inaccessible class of PM-associated and polarized proteins, substantially expanding the molecular landscape of plant cell polarity (**Figure S2**).

The current datasets do not yet provide a complete census of lateral polarity proteins. Several established polar proteins, including BOR1, NIP5;1, and KOIN, were not recovered as ILPP or OLPP candidates, whereas a substantial fraction of the IRK proxitome overlapped with the interactomes of LPC components (**Figure 4E**). Thus, the resulting proxitomes reflect both spatial proximity and physical association, producing a biologically informative but interaction-biased view of lateral PM organization. Further refinement of labeling geometry, expression level, and control design will be required to achieve more comprehensive coverage. Nevertheless, this resource provides a decisive foundation for identifying the molecular determinants of lateral polarity. Comparative analysis of the shared structural, biochemical, and spatial features of these proteins should reveal general principles that specify polarized localization along the radial axis.

Intriguingly, some of the proteins in the IRK proxitome were independently identified as SOSEKI (SOK)-interacting proteins in the accompanying study (*Pukhovaya et al., accompanying manuscript*). Pukhovaya *et al.* identified and designated NRL2, NRL16, and NRL29 as SOSEKI-INTERACTING BTB 1 (SIB1), SIB2, and SIB3, respectively, and AT1G12460 and AT1G62950 as SIB1/NRL2-INTERACTING TRANSMEMBRANE KINASE 1 (SITK1) and SITK2, respectively. They further showed that AT2G40980 exhibited a polarity pattern similar to that of NRL2 and designated it SIB1/NRL2-INTERACTING CYTOSOLIC KINASE (SICK). For clarity, we used the names SITK1, SITK2, and SICK throughout this study. The independent identification of the same set of proteins by different groups, using complementary proteomic approaches combining proximity labeling and co-IP with distinct polar proteins, strongly supports the robustness of these associations and indicates that these proteins constitute a common polarity-associated system in plants.

The LPC is organized around plant-specific NRL scaffold proteins and comprises PATs, SITK1/2, SICK, and IRKI1 (**Figures 2**, **4, and S17**). In addition to NRLs, SICK and IRKI1 are plant-specific proteins (**Figures S17A and S17D**), but their molecular functions remain poorly understood. Notably, IRKI1 contains extensive IDRs (**Figure S17F**), indicating a potential role in mediating multivalent interactions within the LPC together with IDRs of the group C-PATs. Further genetic and biochemical analyses will be required to define how IRKI1 contributes to LPC organization and function.

Upon overexpression, co-expression of PAT19 and NRL2 induced large PM microdomains, and these domains occupied regions largely complementary to CMTs. These observations raise the possibility that LPC microdomains contribute to the establishment or maintenance of lateral PM identity. Consistent with an important developmental role for the LPC, genetic disruption of its components, particularly higher-order mutations in NPT-clade NRLs, caused severe defects in cell morphology in the root meristem. Although the mechanisms linking LPC organization to cell-shape control remain unresolved, the presence of multiple kinases within the LPC indicates that phosphorylation-dependent signaling may operate within these polar microdomains to coordinate cell polarity and morphogenesis.

In animal cells, *S*-acylation, also known as *S-*palmitoylation, contributes to cell polarization by promoting the membrane targeting and activity of specific polarity-associated proteins^30^. In mammalian epithelial cells, the polarity determinant Scribble is *S*-palmitoylated by ZDHHC7, and this modification is required for its targeting to the PM and for normal apicobasal polarity^31^. During epithelial–mesenchymal transition induced by the transcription factor Snail, the depalmitoylase APT2 promotes Scribble depalmitoylation and its displacement from the PM. Accordingly, inhibition of APT2 restores Scribble *S*-palmitoylation, PM localization, and epithelial polarization^32^. In *Drosophila melanogaster*, Fat-dependent planar cell polarity involves the atypical myosin Dachs, whose cortical localization and activity are controlled by the DHHC-type *S*-acyltransferase Approximated (App). App is enriched at the apical cell cortex and is required for the normal cortical accumulation and function of Dachs, thereby contributing to planar cell polarity^33^. Together with these findings in animal cells, our observation that PAT-dependent *S*-acylation promotes microdomain organization at lateral PM domains points to a notable mechanistic convergence in cell polarity systems, in which lipid modification facilitates the membrane recruitment and higher-order organization of polarity-associated proteins.

How LPC polarity itself is established remains unclear. The extracellular domain of IRK is sufficient to specify its polar localization^11,12,34^, indicating that extracellular spatial information may orient IRK and related LRR-RLK VII receptors. These receptors could in turn bias the recruitment of LPC components, with *S*-acylation and higher-order clustering reinforcing the resulting polarized assemblies. Identifying the extracellular information and molecular interactions that initiate this asymmetry will be important for understanding how radial positional information is translated into polarized PM domains.

Finally, the conserved polarity and morphogenetic function of NPT-clade NRLs in Arabidopsis and the liverwort Marchantia indicate that key features of this polarity system arose early in land plant evolution. At the same time, features of LPC organization—including protein clustering, PM microdomain formation, and regulation by *S*-acylation—resemble mechanisms used to establish cell polarity in animals, despite the involvement of largely distinct molecular components. Plants may therefore have combined conserved principles of membrane organization with lineage-specific molecular machinery to generate their characteristic cell polarity and body architecture. Thus, our findings place the LPC at the intersection of cell polarity, morphogenesis, and land plant evolution, establishing a foundation for future studies of how polarity systems are built in plants and for defining the conserved principles of cellular polarization across kingdoms.

### Limitation of this study

This study identified key molecular components of the LPC and demonstrated that PAT-dependent *S*-acylation promotes LPC microdomain assembly and proper NRL2 polarity. However, several mechanistic questions remain unresolved. The large labyrinthine LPC structures were most clearly observed following transient overexpression in tobacco, and analyses at endogenous expression levels will be required to define their native organization and dynamics. Although LPC components showed a mutually exclusive distribution with cortical microtubules, the functional relationship between them remains unclear. Direct evidence that PAT19, PAT20, or PAT22 *S*-acylate NRL2 or other LPC components is also needed to establish how *S*-acylation promotes LPC assembly. Moreover, the molecular cues and trafficking mechanisms that specify distinct lateral PM domains remain unknown. Notably, disruption of PAT-dependent *S*-acylation in *Arabidopsis* redirected NRL2 to an aberrant polarity domain rather than simply reducing its membrane association. Understanding the basis of this altered polarity may provide an important clue to how distinct PM domains are specified and maintained. Finally, how LPC organization regulates cell division, anisotropic growth, and cell-wall remodeling remains to be determined. The components identified here provide a foundation for addressing these questions and understanding how lateral polarity controls plant cell morphogenesis.

## Resource availability

## Lead contact

Further information and requests for resources and reagents should be directed to and will be fulfilled by the lead contact, Akira Yoshinari.

## Material availability

Unique materials generated in this study are available from the lead contact upon request. Gateway destination vectors constructed in this study will be deposited to Addgene.

## Data and code availability

Mass-spectrometry datasets will be deposited to ProteomeXchange Consortium via jPOST. Phylogenetic tree data will be available at iTOL.

## Acknowledgements

We thank D. Weijers for providing materials, engaging in thoughtful discussions, and coordinating the simultaneous submission of the accompanying study. We also thank S. Nasu, K. Hirano, Y. Kawase, K. Sekiguchi, K. Ota, K. Kano, S. Sato, A. Matsumoto, A. Saito, S. Ideguchi, N. Yagi, T. Yamazaki, and A. Tomita for their excellent technical assistance and support. We also thank Y. Mizuta, M. Yamada, N. Uchida, K. Takiguchi, M. M. Wudick, R. Simon, G. Grossmann, G. Vert, K. Torii, K. Nishitani, and J. Takano for helpful discussions and advice. We are grateful to H. Tsutsui, S. Segami, S. Fujita, Y. Sugiyama, T. Sasaki, Y. Oda, T. Nakagawa, M. Ueda, and M. Furutani for kindly providing plasmid vectors. We thank Y. Zhang and S. Li for providing *pat19;pat20;pat22* seeds. We thank the advisory board members of the JST PRESTO consortium entitled Function and Regulation of Plant Molecules for their valuable advice and the Arabidopsis Biological Resource Center (ABRC) for providing the T-DNA insertion lines. This work was supported by JSPS KAKENHI Grant Number JP22K15139, JST PRESTO Grant Number JPMJPR22D9, and the Foundation of Public Interest of Tatematsu, all awarded to A.Y., and JST FOREST Grant Number JPMJFR2253 awarded to K.M. This work was also partially supported by the World Premier International Research Center Initiative (WPI).

## Author contributions

A.Y. conceptualized and designed the study. A.Y., K.Y., K.O., K.F., K.M., A.T., and M.N. performed the experiments. E.M.-S. performed and supervised the proteomic analyses. A.Y. and R.I. conducted the phylogenetic analyses. J.J.L. generated the multisite Gateway vectors. K.Y., K.F., and S.N. generated the Marchantia mutants and transgenic plants. A.Y. and K.O. performed the Y2H assays. A.Y., M.N., S.N., and W.B.F. wrote the manuscript.

## Declaration of interests

The authors declare no competing interests.

## Declaration of generative AI and AI-assisted technologies

During the preparation of this work, the authors used OpenAI ChatGPT (GPT-5.5 and GPT-5.6) for grammar checking and editing. After using this, the authors reviewed and edited the content as needed and take full responsibility for the content of the published article.

## Materials and Methods

### Plant Materials and Growth Conditions

*Arabidopsis thaliana* (L.) Heynh. Columbia-0 (Col-0) ecotype was used as wild type throughout the study. *Nicotiana benthamiana* was used for tobacco agroinfiltration assays. *Marchantia polymorpha* Tak-1 ecotype was used as wild type. The *pat19;pat20;pat22* triple mutant has been characterized previously^23^. The *pSICK:SICK-tdTomato* transgenic line was generated and characterized in the accompanying study (*Pukhovaya et al., accompanying manuscript*). All other transgenic lines and mutants were generated and characterized in this study. Unless otherwise indicated, T_2_-generation transgenic plants were used for experiments. The transgenic lines expressing the PM marker YFP-LTI6b and the nuclear marker H2B-RFP have been described previously ^39,40^.

For transformation of *Arabidopsis thaliana*, the floral dip method using Agrobacterium (*Rhizobium radiobacter*) GV3101 strain carrying a helper plasmid pMP90 was performed as described previously ^41,42^. Transformed plants were selected on half-strength Murashige Skoog (MS) media containing selective antibiotics (20 mg/L hygromycin or 20 mg/L kanamycin) and 50 mg/L claforan or 50 mg/L carbenicillin for prevention of Agrobacterium propagation. For T_1_ selection of CRISPR/Cas9-genome edited lines, red fluorescence of seed coat was used as an indicator.

For transformation of *Marchantia polymorpha,* regenerating thalli were used as described previously, with minor modifications ^43,44^. Apical regions were removed from 14-day-old thalli, and the remaining tissues were cut into pieces and cultured for 3 days to induce regeneration. The regenerating thalli were co-cultivated with Agrobacterium tumefaciens harboring the appropriate binary plasmids in 0M51C medium [Gamborg B5 medium containing 2% (w/v) sucrose and 100 μM acetosyringone] at 22°C for 3 days with agitation at 130 rpm. After co-cultivation, the thalli were washed and transferred to half-strength Gamborg’s B5 agar medium containing hygromycin and cefotaxime. Hygromycin-resistant plants were isolated and propagated for further analyses.

Arabidopsis T-DNA insertion lines were obtained from the Arabidopsis Biological Resource Center (ABRC). SALK_209978 (*nrl2-1*), SALKseq_111979.1 (*nrl2-2*), SALK_040283 (*nrl16-1*), SALK_143045 (*nrl16-2*), GK-860B06 (*nrl19-1*), and SALK_002904 (*nrl29-1*) were used in this study. Genotyping was performed by PCR using specific primers described in **Table S7**. The *nrl2-2;nrl16-2;nrl29-1* (tKO) was generated by crossing. The *nrl2;nrl16;nrl19;nrl24;nrl29;NRL22^Δ9aa^*(qKO) was generated using CRISPR/Cas9 by introducing KMd679 and KMd680 by floral-dip methods with Agrobacterium GV3101 with pMP90. Initially, we attempted to generate a sextuple knockout mutant; however, complete disruption of *NRL22*, which is predominantly expressed in reproductive organs, caused a marked reduction in fertility. We therefore used a line carrying five null mutations together with an *NRL22* allele encoding a nine-amino-acid deletion, which retained sufficient fertility to produce enough seeds for subsequent analyses. The qKO carries the transgenes encoding Sp*Cas9* and the guide RNAs. Genotyping of T-DNA insertion was performed by PCR using KOD FX Neo (Toyobo, Cat#: KFX-201) or QuickTaq HS DyeMix (Toyobo, Cat#: DTM-101) and primers listed in **Table S7**. Genotyping of genome-edited sequences was performed by PCR using KOD FX Neo followed by Sanger sequencing using primers listed in **Table S7**.

For Arabidopsis, seeds were surface-sterilized with 70% (v/v) ethanol, washed with sterile ultrapure water (Milli-Q) four times, and sown on modified MGRL medium^45^ containing 30 µM boric acid, 1% (w/v) sucrose, and 1.5% (w/v) gellan gum in sterile plastic dish (Eiken Chemical, Cat#: LPH-411PFDT-S). The medium plates were incubated at 4°C for 2–3 days and placed vertically in a plant growth chamber (NK System, Cat#: LPH-411PFDT-S) at 22°C with 16 h of light at 60–90 µmol m^−2^ s^−1^ and 8 h of dark. For Marchantia, gemmalings were put on half-strength Gamborg B5 medium containing 1.2% (w/v) agar or MGRL medium containing 30 µM boric acid, solidified with 1.5% (w/v) gellan gum. The medium plates were incubated in a plant growth chamber (NK System, Cat#: LPH-411SP) at 22°C with 16 h of light at 60–90 µmol m^−2^ s^−1^ and 8 h of dark or constant light at 30–50 µmol m^−2^ s^−1^. For tobacco, plants were grown in plant growth room at 22°C with 16 h of light at 60–80 µmol m^−2^ s^−1^.

### Plasmid construction

All plasmids and oligonucleotide identifiers used in this study are listed in **Tables S7–S10**. DNA fragments were amplified using PrimeSTAR GXL (Takara Bio, Cat#: R050A), KOD One (Toyobo, Cat#: KMM-201), or Phusion High-Fidelity DNA Polymerase (Thermo Fisher Scientific, Cat#: F530). *Arabidopsis thaliana* Col-0 genomic DNA and a Col-0 cDNA library prepared from our lab stock, or existing plasmids were used as PCR templates. Unless otherwise indicated, coding and genomic fragments intended for C-terminal fusion were amplified without their native stop codons.

To generate pRPS5A:GW:mYFP (pAY772), the *RPS5A* promoter was amplified from pKAMA-ITACHI1.1R ^46^ using primers 1743 and 3180 and assembled with HindIII/XbaI-digested pGWB441m by Gibson Assembly® Cloning Kit (NEB, Cat#: E2611S). The *HSP* terminator-containing derivative pRPS5A:GW:mYFP (pAY767) was subsequently generated by Gibson assembly using a fragment amplified with primers 3206 and 3207 and a pAY772-derived backbone amplified with primers 3208 and 3209. To generate the pSCR:GW:mYFP destination vector pAY889, pGWB441m was digested with HindIII and XbaI, and a PCR-amplified *SCR* promoter fragment was inserted using In-Fusion Cloning Kit (Takara Bio, Cat#: 639648). The pUBQ10:GW:mGFP vector pAY187 was generated by introducing the *UBQ10* promoter into HindIII/XhoI-digested pGWB504m by restriction–ligation. To prepare destination vectors carrying fluorescent proteins followed by the HSP terminator, a DNA fragment containing XbaI, ApaI, KpnI, and EcoRV restriction sites, *mCerulean*, and the *HSP* terminator was synthesized by Integrated DNA Technologies (IDT) (**Table S8**) and introduced into AscI/SalI-digested pGWB504m to generate pAY1740. The pRPS5A:mYFP:GW vector pAY1042 was generated by assembling the *mYFP:GW (attR1-Cm^R^-ccdB-attR2)* sequence with an inverse PCR-amplified pAY767 backbone using NEBuilder HiFi. The *mYFP:GW* sequence was amplified from pGWB442m using primers 4256 and 4257, and the pAY767 backbone was amplified by inverse PCR using primers 4258 and 4259. The *mCherry*-containing vector pAY1778 was generated by inserting an amplified *mCherry* fragment into XbaI/EcoRV-digested pAY1740 using NEBuilder® HiFi DNA Assembly (NEB, Cat#: E2621S). The corresponding monomeric Green Fluorescent Protein (mGFP) vector pAY1779 was generated by inserting an *mGFP* fragment with primers 7361 and 7363 into XbaI/EcoRI-digested pAY1740 by NEBuilder® HiFi DNA Assembly. The pUBQ10:GW:mCherry vector pAY1782 was generated by ligating an *mCherry-HSP* terminator fragment derived from pAY1778 into EcoRI/AscI-digested pAY187. Similarly, pUBQ10:GW:mGFP (pAY1783) was generated by ligating the *UBQ10* promoter fragment obtained from pAY187 into EcoRI/AscI-digested pAY1779. The StayGold^E138D^ (mSG) fragment was amplified from pcDNA3-10His-mStayGold (E138D)^47^ using primers 7382 and 7383 and used to generate the no-promoter Gateway vector pAY1825. The pUBQ10:GW:StayGold^E138D^(mSG) vector pAY1826 was subsequently generated by ligating the *UBQ10* promoter fragment obtained from pAY187 into EcoRI/AscI-digested pAY1825.

To generate pENTR P4-P1r-pSCR, a 2,000-bp fragment upstream of the *SCR* start codon was amplified from *Arabidopsis thaliana* Col-0 genomic DNA using primers p4p1SCR_F and p4p1SCR_R. The PCR product was inserted into BamHI/XhoI-digested pENTR P4-P1r using In-Fusion cloning (Takara Bio). To generate pENTR P2r-P3-V5-TurboID-GFP-stop-terminator, a GFP fragment was amplified from synthesized DNA (**Table S8**) using primers p2p3GFP_F and p2p3GFP_R. The PCR product was inserted into BamHI/KpnI-digested pENTR P2r-P3-stop-terminator ^48^ using In-Fusion cloning, generating pENTR P2r-P3-GFP-stop-terminator. The resulting plasmid was subsequently digested with AscI, and a synthesized V5-TurboID fragment (**Table S8**) was inserted using In-Fusion cloning.

The *CaMRLK/MEE62* coding sequence was amplified from a Col-0 cDNA library using primers 1599 and 1600 and cloned into pENTR/D-TOPO (Thermo Fisher Scientific, Cat#: K240020SP) to generate pAY416. An *IRK* genomic fragment containing introns was amplified from Col-0 genomic DNA using primers 3384 and 3385 and cloned into pENTR/D-TOPO, generating pAY837. Because pAY837 contained a mutation, the wild-type IRK entry clone pAY861 was subsequently generated by inverse amplification using primers 3477 and 3476, followed by NEBuilder HiFi assembly. A *GUS-GFP* fragment was amplified using primers 3719 and 3720 and cloned into pENTR/D-TOPO to generate pAY960. Genomic fragments of *UCHIGAWA* (AT2G23700), *SCAIL1* (AT4G40050), *PLPDT1* (AT2G23520), *PLPDT2* (AT4G37100), *NRL2*, *NRL16*, *NRL19*, *NRL22*, *NRL24*, *NRL29*, *PAT19*, *PAT20*, *PAT22*, *SITK1*, *SITK2*, and *IRKI1* were amplified from Col-0 genomic DNA and introduced into pDONR221. The corresponding primer pairs are listed in **Table S7**. ORF entry clones for *NRL2*, *NRL16*, *NRL19*, *NRL28*, *NRL29*, *PAT19*, *PAT22*, and *IRKI1* were generated by inverse PCR of the corresponding genomic entry clones by removing their promoter sequence or amplifying the ORF sequences from *A. thaliana* cDNA library as templates. pDONR221 entry clones were generated for *PBL35*, *PSI3* (AT5G08660), *PSIL1* (AT3G23160), *PSIL3* (AT5G04550), *PP2A B′θ*, *RLCK-VI B1*, *MSL4*, and *SCAR1*/*WAVE1*. The pENTR/D-TOPO entry clone carrying the AT3G47210 ORF without a stop codon was generated by PCR using primers 3967 and 3968, followed by a TOPO cloning reaction. Because the C-terminal mGFP fusion of AT3G47210 did not produce detectable fluorescence, a stop codon was introduced into the AT3G47210 entry clone (pAY1023) by site-directed mutagenesis using primers 5544 and 5546. The resulting entry clone was recombined with pAY1042 using LR Clonase II. The amplified fragments were introduced into pDONR221 using BP Clonase II Enzyme Mix (Thermo Fisher Scientific, Cat#: 11789020). Because the original *NRL2* and *NRL16* genomic entry clones contained mutations, wild-type *NRL2* and *NRL16* entry clones were generated by site-directed mutagenesis using primers 4825 and 4826 for *NRL2* and primers 4827 and 4828 for *NRL16*, respectively. The pDONR221/mCherry-HSPter entry clone pAY1836 was generated by amplifying the mCherry-HSPter fragment from pAY1778 using PrimeSTAR GXL DNA polymerase and primers 7397 and 7398, followed by BP recombination into pDONR221.

For construction of the Mp*NPT* expression vector in *Arabidopsis thaliana*, the Mp*NPT* coding sequence was amplified from a *Marchantia polymorpha* Tak-1 cDNA library using primers 6974 and 7431 and cloned into pDONR221 by BP recombination. Because the resulting entry clone contained an unexpected sequence within the CDS, the sequence was removed by inverse PCR using primers 7707 and 7708. The PCR product was then circularized using the NEBuilder HiFi DNA Assembly system, yielding the corrected entry clone pAY2149.

Binary expression constructs were generated by Gateway LR recombination using LR Clonase II Enzyme Mix (Thermo Fisher Scientific) between the appropriate entry and destination vectors listed in **Table S9**. These included *pRPS5A*-, *pSCR*-, *p35S*-, or *pUBQ10*-driven constructs expressing *CaMRLK*, *BRI1*, *NRL*s, *PAT*s, *SITK1*/2, *IRKI1*, *PBL35*, and the other candidates as C-terminal fusions with *mGFP*, *mYFP*, *mCherry*, *StayGold^E138D^* (*mSG*), *mRFP*, or *GUS*. Genomic mGFP fusion constructs were generated for *UCHIGAWA*, *SCAIL1*, *PLPDT1*/2, *NRL2*, *NRL16*, NRL19, *NRL29*, *PAT19*, *PAT20*, *PAT22*, *SITK1/2*, *IRKI1*, *PSI3*, *PSIL1*, *PSIL3, PP2A B′θ*, *RLCK-VI B1*, *MSL4*, and *SCAR1*. Genomic GUS fusion constructs for *NRL2*, *NRL16*, *NRL19*, *NRL22*, *NRL24*, and *NRL29* were generated by Gateway LR recombination using pGWB533^49^ as the destination vector. Ectopic and heterologous expression constructs were generated for *PAT22*, *PSIL1*, *PSIL3*, *NRL28*, *MAB4*, Mp*NPT,* and selected LPC components using constructs driven by the *RPS5A*-, *SCR*-, or *UBQ10* promoters. The pSCR:PBL35-mYFP construct pAY1357 was generated by Gateway LR recombination between the *PBL35* entry clone pAY1315 and the pSCR destination vector pAY889. The pUBQ10:mScarlet-HDEL construct pAY2200 was generated by Gateway LR recombination between the entry clone opSF356, kindly provided by Dr. S. Fujita, and the destination vector pGWB4-UBQ10, kindly provided by Dr. S. Segami.

The pSCR:CaMRLK-TurboID-GFP (pAY591), pSCR:IRK-TurboID-GFP (pAY898), and pSCR:GUS-GFP-TurboID-GFP (pAY991) constructs were generated by Multisite Gateway recombination using LR Clonase II Plus (Thermo Fisher Scientific, Cat#: 11791020). The reactions combined pENTR P4-P1r-pSCR, the corresponding middle entry clone—pAY416, pAY861, or pAY960—and pENTR P2r-P3-V5-TurboID-GFP-stop-terminator with the pHm43GW destination vector.

Three PAT19 C-terminal truncation entry clones, pAY1657, pAY1658, and pAY1659, were amplified from the PAT19 ORF entry clone pAY1456 using the common forward primer 6740 and reverse primers 6741, 6742, and 6743, respectively. The three constructs encoded PAT19Δ389–718, PAT19Δ609–718, and PAT19Δ651–718, respectively. The resulting PAT19 truncation variants were transferred into the pUBQ10:GW:mGFP destination vector by Gateway LR recombination to generate pAY1764, pAY1765, and pAY1766.

For Y2H analyses, full-length NRL2 and the PAT19 C-terminal region were amplified and introduced into pGBKT7 to generate pAY1975 and pAY1976, respectively. NRL2, NRL16, NRL19, NRL28, NRL29, MAB4, SICK, and the kinase domains of SITK1, SITK2, and IRK were amplified and introduced into pGADT7, generating pAY1977–pAY1979, pAY1982–pAY1984, and pAY1987–pAY1990. The amplified inserts were assembled with the corresponding linearized pGBKT7 or pGADT7 backbones using either ligation or NEBuilder HiFi DNA Assembly.

For the DNA construction of MpNPT-Citrine, a genomic fragment of *MpNPT* lacking the stop codon was amplified using PrimeSTAR GXL DNA Polymerase with the primers pENMpNPTFor and pENMpNPTnsRev. A 5-kb region upstream of MpNPT was amplified using the primers proMpNPT_for and proMpNPT_Rev. The amplified MpNPT and promoter fragments were inserted into pENTR/D-TOPO and pDONR P4-P1R, respectively, using the In-Fusion cloning system. To generate proMpNPT:MpNPT-mCitrine, the promoter entry clone (pDONR_proMpNPT) and the MpNPT entry clone (pENTR_MpNPT) were recombined with R4pMpGWB339 using a MultiSite Gateway LR reaction^50^. For constitutive expression, pENTR_MpNPT was recombined with pMpGWB306 or pMpGWB308 using Gateway LR reactions to generate pro35S:MpNPT-Citrine and proMpEF1α:MpNPT-Citrine, respectively ^51^.

To generate the CRISPR/Cas9 sgRNA expression construct, pMpGE010_MpNPTg1 for Marchantia, pMpGE_En01 was linearized by digestion with SacI and PstI. Complementary oligonucleotides, MpNPT_Crispr1For and MpNPT_Crispr1Rev, were annealed to generate double-stranded DNA fragment containing the guide RNA sequence, which was introduced into the linearized pMpGE_En01 vector using the In-Fusion cloning method. The resulting entry clone was subsequently recombined with pMpGE010 by a Gateway LR reaction. The oligonucleotide sequences are listed in **Table S7**.

To generate the CRISPR/Cas9 sgRNA expression constructs KMd679 and KMd680, the pVS1 rep region of pMDC123 ^52^, kindly provided by Dr. M. Ueda, was amplified using primers KMol463 and KMol464 and ligated into the BsaI/NheI-digested pMDC123 backbone, thereby removing the endogenous BsaI site. A fragment containing the RPS5A promoter and a ccdB cassette flanked by BsaI sites was then amplified from pWAT235-hcoCas9x6-cc using primers KMol478 and KMol479, whereas the modified pMDC123 backbone was amplified using primers KMol480 and KMol481. The two fragments were assembled using seamless ligation cloning extract (SLiCE) ^53^. A Cas9io-containing fragment was subsequently amplified from pRU295 ^54^ using primers KMol1049 and KMol1050 and inserted into the BcuI/BamHI-digested intermediate vector using SLiCE. Finally, a pAt2S3-mCherry-KDEL fragment amplified from KMd104 ^55^ using primers KMol493 and KMol494 or a synthesized pAt2S3-EGFP-KDEL fragment (**Table S8**) was inserted into the XbaI-digested vector using NEBuilder® HiFi DNA Assembly, generating geM056 and geM050, respectively. sgRNA shuttle vectors were derived from pBluescript II SK(+) (Stratagene), in which an endogenous BsaI recognition site was disrupted by site-directed mutagenesis using primers pBS-dBsa-F and pBS-dBsa-R. Six Arabidopsis U6 fragments (U6-26P, U6-6P, U6-29P, U6-4P, U6-5P, and U6-6P) amplified from Col-0 genomic DNA and an sgRNA scaffold derived from pUC119-gRNA ^56^ were used to construct the six corresponding shuttle vectors, into which a *ccdB* cassette was subsequently introduced. The sgRNA scaffold was further replaced with the M4-5bp modified scaffold ^57^, generating pBSx1-pU626-M4-5bp-cc, pBSx2-pU66-M4-5bp-cc, pBSx3-pU629-M4-5bp-cc, pBSx4-pU64-M4-5bp-cc, pBSx5-pU65-M4-5bp-cc, and pBSx6-pU66-M4-5bp-cc. Complementary oligonucleotides encoding guide sequences targeting NRL2, NRL16, NRL19, NRL22, NRL24, and NRL29 were annealed and introduced into the corresponding shuttle vectors by BsmBI-based Golden Gate cloning using BsmBI (NEB, Cat#: R0580) and T7 DNA Ligase (NEB, Cat#: M0318S). The six resulting sgRNA expression cassettes were assembled into geM050 and geM056 by BsaI-based Golden Gate cloning using BsaI-HF v2 (NEB, Cat#: R3733) and T7 DNA Ligase, yielding KMd679 and KMd680, respectively. The oligonucleotide sequences are listed in **Table S7**.

### TurboID-based proximity labeling

Roots of 9-day-old Arabidopsis seedlings grown vertically on MGRL medium plates containing 1 µM boric acid, 1% (w/v) sucrose, and 1.5% (w/v) gellan gum were excised and transferred into 50 mL MGRL liquid medium containing 1 µM boric acid, 0.5% (w/v) sucrose, and 50 µM biotin and incubated for 3 h at room temperature without shaking. Then, roots were washed three times with ice-cold deionized water (Elix). After washing, excess surface water was gently removed from the roots using paper towels, and the fresh weight of each sample was measured. The root samples were then transferred to 10 mL Multi-Beads Shocker (MBS) tubes (Yasui Kikai, Osaka, Japan) containing metal cones, immediately frozen in liquid nitrogen, and stored at −80°C until protein extraction.

### Protein extraction and concentration of biotinylated proteins for mass-spec analysis

Frozen root samples were disrupted using a Multi-Beads Shocker MB3000 (Yasui Kikai) at 1,500 rpm for 30 s, repeated three times with liquid nitrogen cooling between cycles. After incubation at −30°C for 30 min, proteins were extracted by adding modified SDT buffer [125 mM Tris-HCl (pH 7.5), 5%(w/v) sodium dodecyl sulfate (SDS), 0.125 M dithiothreitol (DTT)]^58^ at 0.7 mL/g fresh weight, followed by disruption at 1,000 rpm for 10 s. Extracts were boiled for 5 min and centrifuged twice at 15,000 rpm for 15 min to remove debris. Protein concentrations were measured, and 1 mg protein was adjusted to 500 µL.

Free biotin was removed using PD-10 desalting columns (Cytiva, Cat#: 52130800) equilibrated with 1/10 SDT buffer. The 1 mg protein sample was diluted to 2.5 mL with Milli-Q ultrapure water and loaded onto the column. Proteins were eluted with 3.5 mL of 1/10 SDT buffer. 500 µg of total protein was diluted with dilution buffer [50 mM Tris-HCl (pH 7.5), 150 mM NaCl, 0.2% (v/v) Triton X-100], and mixed with 100 µL µMACS Streptavidin microbeads (Miltenyi Biotech, Cat#: 130-133-282), and incubated overnight at room temperature with gentle rotation.

Biotinylated proteins were purified using µMACS columns and separators (Miltenyi Biotech, Cat#: 130-042-602, 130-042-701). Columns were equilibrated with 100 µL equilibration buffer supplied by the manufacturer, followed by two washes with 100 µL PBS containing 0.5% (w/v) SDS. The bead-bound protein solution was then applied to the column. Columns were washed sequentially with PBS, 1 M KCl, 0.1 M Na_2_CO_3_, and 2 M urea in 10 mM Tris-HCl (pH 8.0), and the columns were immediately subjected to on-column digestion.

### RNA extraction and cDNA synthesis

Total RNA was extracted from the thalli of *Marchantia polymorpha* (Tak-1) using the RNeasy Plant Mini Kit (QIAGEN, Cat#: 74904) according to the manufacturer’s instructions. For cDNA library preparation, 400 ng of total RNA was subjected to reverse transcription using the ReverTra Ace qPCR RT Master Mix with gDNA Remover (TOYOBO, Cat#: FSQ-301). Briefly, the RNA template was treated with 4x DN Master Mix at 37°C for 5 min to remove contaminating genomic DNA. The reverse transcription reaction was then performed with 5x RT Master Mix II under the following conditions: incubation at 37°C for 15 min, followed by 50°C for 5 min for extension, and heat inactivation of the enzyme at 98°C for 2 min. The resulting cDNA library was used as a template for PCR amplification of MpNPT.

### Confocal laser scanning microscopy

A Zeiss LSM800 equipped with 40x water-immersion (1.1 NA LD C-Apochromat, Zeiss) and 63x oil-immersion (1.4 NA Plan-Apochromat, Zeiss) objective lenses and 405-, 488-, and 561-nm lasers and a Leica TCS-SP8 gSTED, equipped with 63x oil-immersion (1.40 NA HC PL APO CS2, Leica) objective lens and a white-light laser, were used for confocal imaging. For the staining of cell walls in living cells, plants were incubated in deionized water (Elix water; Merck Millipore) containing 10 µg/mL propidium iodide (PI) (Fujifilm Wako Pure Chemical, Cat#: 169-26281) for 5 – 30 min. For the staining of PM in living cells, plants were incubated in MGRL liquid medium containing 30 µM boric acid and 1 µM FM4-64 (Thermo Fisher Scientific, Cat#: T3166) for 0.5 – 1 h. For LSM800, excitation/detection wavelengths were 405/410–470 nm for SR2200 dye, 488/505–545 nm for GFP, mGFP, YFP, mYFP, and mSG, 561/576–700 nm for mRFP, mCherry, tdTomato, mScarlet, and PI dye, 488/650–710 nm for FM4-64 dye. For TCS-SP8, excitation/detection wavelengths were 488/500–530 nm for mGFP and 561/580–650 nm for mCherry.

For the fixation of *M. polymorpha* gemmalings expressing MpNPT-mCitrine, plants were incubated with 4% (w/v) paraformaldehyde in PBS for 1 h at room temperature under vacuum. After washing with PBS, the samples were cleared with ClearSee solution as described previously ^44,59^ and mounted in ClearSee solution. Fluorescence was observed using a Nikon AX R confocal laser scanning microscope equipped with 40x water-immersion (1.15 NA LWD Lambda S 40XC WI, Nikon). Excitation/detection wavelengths were 488/502–546 nm.

### 3D cell segmentation and volume analysis

Three-dimensional confocal image stacks of 2-day-old gemmalings stained with SR2200 were analyzed using MorphoGraphX ver. 2.0.3. Individual cells were segmented using the ITK Watershed Auto Seeded algorithm. The segmented images were converted into three-dimensional cell meshes using the Marching Cubes 3D process, and cell volumes were quantified using Cell Analysis 3D.

### Co-immunoprecipitation for proteomic analysis

Plants were grown vertically for 14 days on MGRL medium plates containing 30 µM boric acid, 1% (w/v) sucrose, and 1.5% (w/v) gellan gum. Roots were excised, weighed, transferred to 5-mL MBS tubes, and immediately frozen in liquid nitrogen. The frozen samples were pulverized using a Multi-Beads Shocker at 2,000 rpm for 30 s, repeated three times. The resulting frozen powder was transferred to a pre-chilled mortar containing ice-cold extraction buffer [50 mM Tris-HCl (pH7.5), 100 mM NaCl, 10% (v/v) glycerol, 1 mM ethylenediaminetetraacetic acid (EDTA), 5 mM DTT, cOmplete Mini EDTA-free (Merck Millipore, Cat#: 4693159001), 1 mM Pefabloc SC (Merck Millipore, Cat#: 11429868001), 2% (v/v) Nonidet P-40] at a ratio of 3 mL per gram fresh weight and further homogenized with a pestle. The homogenate was centrifuged once at 12,000 rpm for 15 min at 4°C to remove large debris. The resulting supernatant was further clarified through a 0.22 µm cellulose acetate (CA) syringe filter (Membrane Solutions, Cat#: CA013022) in a cold room. Protein concentrations were determined using the Pierce™ 660 nm Protein Assay with Ionic Detergent Compatibility Reagent (Thermo Fisher Scientific, Cat#: 22662, 22663) and a Spark Microplate Reader (Tecan). The samples were then adjusted to contain 800 µg of total protein in a final volume of 1,650 µL by diluting with the extraction buffer.

For affinity purification, 50 µL of µMACS Anti-GFP magnetic beads (Miltenyi Biotech, Cat#: 130-091-125) was added to each protein extract, and the samples were incubated on a rotator for 1 h. The samples were subsequently applied to µMACS columns. Each column was washed twice with 200 µL of extraction buffer, four times with 200 µL of wash buffer 1 supplied by the manufacturer, and three times with 200 µL of 50 mM Tris-HCl (pH 7.5). A low-protein-binding collection tube (Watson, Cat#: PK-15C-500N) was then placed beneath each column for subsequent sample collection.

### On-column digestion with trypsin and lysylendopeptidase C

For on-column digestion, 25 µL elution buffer I [1.94 M urea, 48.5 mM Tris-HCl (pH 7.5), 5 ng/µL Lys-C, and 5 ng/µL trypsin, 1 mM DTT] was added to each column and incubated for 1 h at room temperature for partial digestion. Peptides were then eluted twice with 50 µL elution buffer II [1.98 M urea, 49.5 mM Tris-HCl (pH 7.5), 5 mM CAA]. The eluates were collected into protein low-binding tubes and further digested for 20 h at 26°C with gentle shaking.

### LC-MS/MS analysis

Peptides were analyzed using an Orbitrap Astral mass spectrometer (Thermo Fisher Scientific) coupled to a Vanquish Neo nanoLC system (Thermo Fisher Scientific) equipped with a nano-electrospray ionization source. Peptides were separated on a reversed-phase NTCC-360/75-1.9-123 C18 column (75 μm i.d. × 120 mm, 1.9 μm; Nikkyo Technos Co., Ltd.) at a flow rate of 300 nL/min maintained at 50°C using a gradient of solvent B (0.1% formic acid in 99.9% acetonitrile) in solvent A (0.1% formic acid in water).

For data-independent acquisition (DIA), full MS scans were acquired in the Orbitrap analyzer over an m/z range of 380–980 followed by DIA MS/MS scans acquired in the Astral analyzer using 300 variable isolation windows covering the precursor mass range. The normalized collision energy (NCE) was set to 25. The automatic gain control (AGC) target and maximum injection time were controlled automatically by the instrument. The total cycle time was approximately 2 – 3 s, providing sufficient sampling across chromatographic peaks.

An *in silico* spectral library was generated from the Arabidopsis thaliana TAIR10 protein database using DIA-NN (ver. 2.2.0), and raw DIA data were analyzed in library-free mode. Carbamidomethylation of cysteine was specified as a fixed modification, while oxidation of methionine and protein N-terminal acetylation were set as variable modifications. Protein inference and quantification were performed using the default DIA-NN settings. Peptide and protein identifications were filtered at a false discovery rate (FDR) of 1% at both the precursor and protein levels.

### Statistical analysis of proteome data

Perseus version 3.0.0.0 (MaxQuant) was used for statistical analysis of the proteomic data. From the available biological replicates, three replicates with comparable numbers of identified proteins were selected for each condition. Protein entries with missing values in more than 10 of the 12 samples were removed using the “Filter rows based on valid values” function. The remaining missing values were imputed from a normal distribution.

### Phylogenetic analysis

Protein sequences of five *Arabidopsis thaliana* genes were independently used as queries to identify homologous proteins across representative green plant lineages. Protein datasets were obtained from publicly available genome resources, including JGI Phytozome, Ensembl Plants, NCBI RefSeq, and species-specific genome databases (**Table S5**).

Homolog candidates were identified by BLASTP searches (BLAST+ v2.17.0) against each species-specific protein dataset. Candidate sequences with an E-value ≤ 1 × 10^−20^ and an alignment length of at least 100 amino acids were retained for subsequent analyses. When multiple transcript isoforms were annotated for a gene, the longest protein isoform was selected. Sequence identifiers were standardized prior to multiple sequence alignment.

Multiple sequence alignments were generated using MAFFT v7.526 ^60^ with the --auto option. Poorly aligned regions were removed using trimAl v1.5.1^61^ with the -automated1 option. Maximum-likelihood phylogenetic trees were inferred using IQ-TREE 3 v3.1.3^62^. The best-fitting amino acid substitution model was selected using ModelFinder, and branch support was evaluated with 1,000 ultrafast bootstrap replicates (-m MFP -bb 1000 -nt AUTO). Phylogenetic trees were visualized and annotated using Interactive Tree Of Life (iTOL) v7 ^63^.

For the phylogenetic analysis of SCAIL1, representative fungal homologs and the human SCAI protein were additionally included as reference sequences. These sequences were obtained from NCBI RefSeq and are listed in **Table S6**.

### Agroinfiltration

*Nicotiana benthamiana* plants grown for 2 – 4 weeks under a 16-h light/8-h dark cycle at 22°C were subjected to Agrobacterium-mediated transient gene expression by agroinfiltration. Plants of the same age were used within each experiment. Agrobacterium cells were washed and resuspended in infiltration buffer containing 10 mM MES-NaOH (pH 5.6) and 10 mM MgCl_2_ and infiltrated into the leaves. Imaging was performed 66–72 h after infiltration.

### Morphological analysis of root apical meristem

Three-day-old Arabidopsis seedlings grown on MGRL medium were fixed in 4% (w/v) paraformaldehyde in phosphate buffer for 1 h under vacuum. The specimens were then washed three times with PBS and infiltrated with ClearSee solution containing 10% (w/v) xylitol, 15% (w/v) sodium deoxycholate, and 25% (w/v) urea ^59^. After a 1-day incubation with ClearSee, the cell walls were stained with 0.1% (v/v) SCRI Renaissance Stain 2200 (SR2200) fluorescent dye (Renaissance Chemicals) diluted in ClearSee solution immediately before imaging.

### Histochemical GUS staining and imaging

Plant samples were immersed in 90% (v/v) acetone and incubated for 10 – 15 min for fixation. The acetone was removed, and the samples were rinsed three times with GUS buffer [50 mM sodium phosphate buffer (pH 7.2), 0.5 mM potassium ferricyanide [K_3_Fe(CN)_6_], and 0.5 mM potassium ferrocyanide [K_4_Fe(CN)_6_]. The samples were then completely submerged in the GUS staining solution [GUS buffer containing 1 mM X-gluc] and vacuum-infiltrated until they sank. Samples were incubated at 37°C for 4 h (roots) or overnight (flowers). After staining, the samples were rinsed three times with 70% (v/v) ethanol and incubated in 70% ethanol overnight (roots) or 3 days (flowers). For floral organs, samples were subjected to clearing by immersing in chloral hydrate clearing solution consisting of chloral hydrate, water, and glycerol at a ratio of 8:3:1 for 3 days. Light microscopy of the GUS-stained specimens was performed using a Zeiss AxioImager.A2 equipped with 10x objective (0.45 NA Plan-APOCHROMAT, Zeiss) and Axiocam 512 color (Zeiss). Stereo microscopy of the specimens was performed using an SMZ 18 (Nikon) equipped with WRAYCAM-NO A2000 (Wraymer).

### Yeast two-hybrid (Y2H) assay

Bait and prey plasmids were co-transformed into the Y2HGold yeast strain (Takara Bio, Cat#: 630498). Yeast cells grown overnight in YPDA liquid medium [1% (w/v) Yeast Extract, 2% (w/v) poly-peptone, 2% (w/v) D(+)-glucose, 120 mg/L adenine sulfate] were collected, washed, and resuspended in One-Step solution containing 40% (w/v) polyethylene glycol, 200 mM lithium acetate, 100 mM DTT, 1 mM EDTA, and 10 mM Tris-HCl (pH 7.5), modified from a previous report ^64^. Salmon sperm DNA and plasmid DNA were added to final concentrations of 1 mg mL^−1^ and approximately 15 µg mL^−1^, respectively. The cells were incubated at 42°C for 1 h, diluted fourfold, and spread onto synthetic complete medium lacking Leu and Trp (-LT). Plates were incubated at 30°C for 3 days. For Y2H assays, transformants were cultured overnight in liquid -LT medium. The cultures were serially diluted 10-fold with sterile water, and 10-µL aliquots were spotted onto -LT medium and either SD medium lacking Leu, Trp, and His (-LTH) or Leu, Trp, His, and Ade (-LTHA). Plates were incubated at 30°C for 3–5 days. The p53 bait and SV40 large T-antigen prey plasmids supplied with the Matchmaker Gold Yeast Two-Hybrid System (Takara Bio, Cat#: 630489) were used as positive controls.

### Quantification of polarity index

Polarity index was quantified as described previously ^65^. Briefly, the apical and basal PM domains of individual cells were traced using the segmented-line tool in Fiji, and the fluorescence intensities in the outer and inner halves of each PM domain were quantified. The fluorescence intensity in the outer half was divided by that in the inner half, and the mean value obtained from the apical and basal PM domains was used as the polarity index for each cell. Thus, values greater than 1 indicate outer-lateral polarity, whereas values less than 1 indicate inner-lateral polarity. Statistical analysis was performed by Prism 10 (GraphPad).

### Genome editing of Arabidopsis thaliana and Marchantia polymorpha

Genome editing of the NPT-clade *NRLs* in *Arabidopsis thaliana* was performed by co-introduction of the multiplex CRISPR/Cas9 constructs KMd679 and KMd680. These constructs carry genes encoding a seed storage protein OLE-1 fused with GFP or TagRFP. T_1_ seeds were selected based on the fluorescence intensity of the seeds. Mutations were examined by PCR followed by Sanger sequencing using specific primers listed in **Table S7**. Genome editing of Mp*NPT* (Mp7g03290) in *Marchantia polymorpha* was performed as described previously ^66^. Mutations were examined by PCR followed by Sanger sequencing using specific primers listed in **Table S7**.

### Phenotypic analysis of *Marchantia polymorpha*

For the growth analysis of Marchantia, gemmalings were placed onto MGRL medium containing 1% (w/v) sucrose and 1.5% (w/v) gellan gum for 16 days under 16-h/8-h light/dark conditions at 22°C.

### Prediction of intrinsically disordered regions

The C-terminal tail region predicted by ConPred II ^67^, integrated in aramemnon server (https://aramemnon.botanik.uni-koeln.de/), was subjected to AIUPred (https://aiupred.elte.hu/) ^68^ to predict intrinsically-disordered regions (IDRs).

### Accession numbers

CaMRLK (AT5G45800), IRK (AT3G56370), NRL2 (AT1G30440), NRL16 (AT3G44820), NRL19 (AT1G69430), NRL22 (AT5G25250), NRL24 (AT5G11680), NRL29 (AT5G66560), PAT19 (AT4G15080), PAT20 (AT3G22180), PAT22 (AT1G69420), IRKI1 (AT5G12900), SICK (AT2G40980), SITK2 (AT1G62950), SITK1 (AT1G12460), NRL28 (AT5G48800), MAB4 (AT4G31820), NPH3 (AT5G64330), CKL7 (AT5G44100), TUB6 (AT5G12250), UCHIGAWA (UCG; AT2G23700), SCAIL1 (AT4G40050), PLPDT1 (AT2G23520), PLPDT2 (AT4G37100), SCR (AT3G54220), SHR (AT4G37650), BRI1 (AT4G39400), Lti6b (AT3G05890), PP2A B′θ (AT1G13460), PSI3 (AT5G08660), PSIL1 (AT3G23160), PSIL3 (AT5G04550), PXC2/CANAR (AT5G01890)

## Notes

### Competing Interest Statement

The authors have declared no competing interest.

