## Supplemental Figures for "The Lateral Protein Cluster as a Key Component of Plant Cell Polarity"

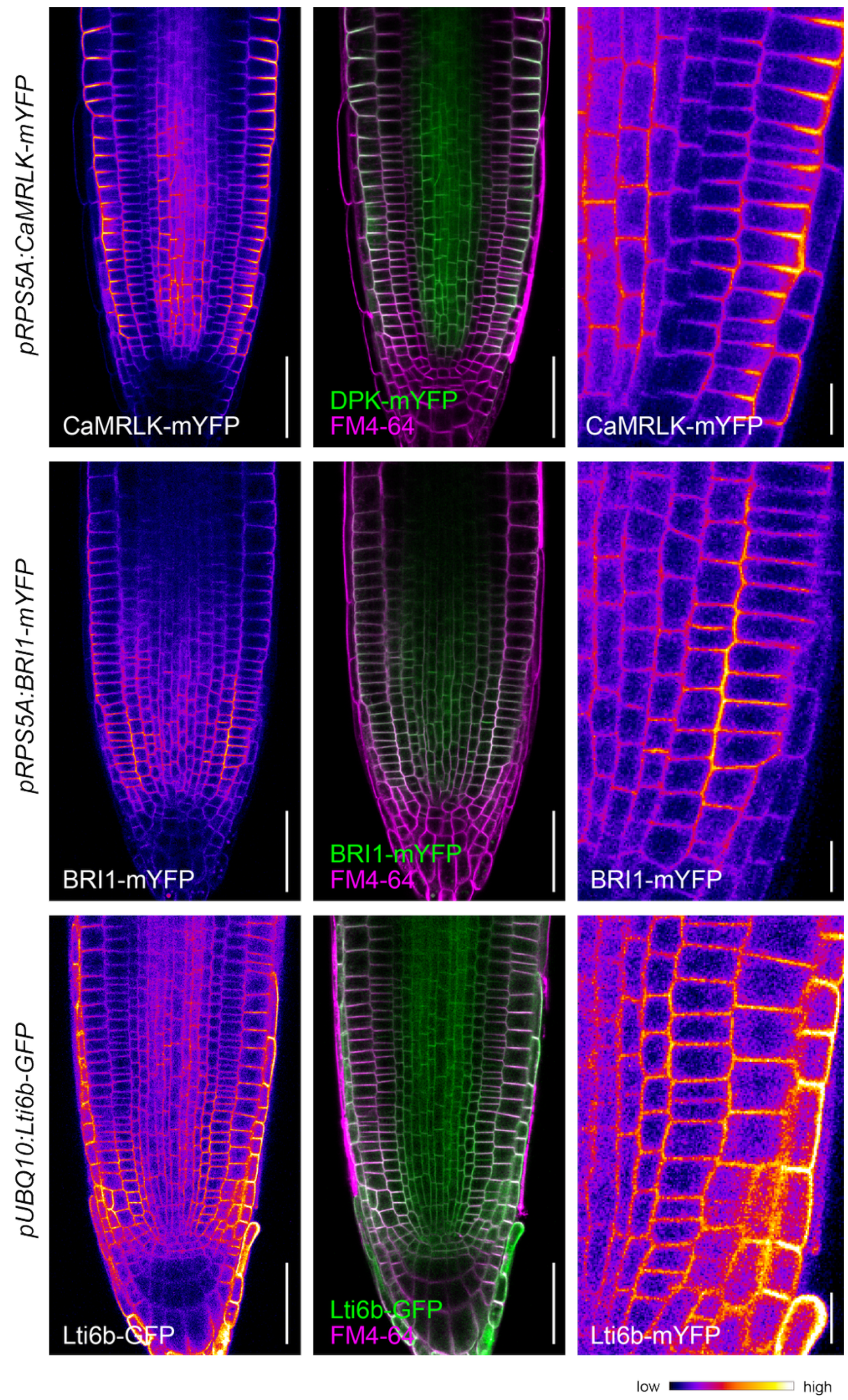


**Figure S1. Subcellular localization of CaMRLK is distinct from those of BRI1 and Lti6b.** Confocal images of primary root tip of transgenic Arabidopsis thaliana plants harboring either *pRPS5A:CaMRLK-mYFP*, *pRPS5A:BRI1-mYFP*, or *pUBQ10:Lti6b-GFP*. Four-day-old seedlings grown on MGRL medium were stained with 1 µM FM4-64. Scale bar: 50 µm (left and middle panels) and 10 µm (right panels).

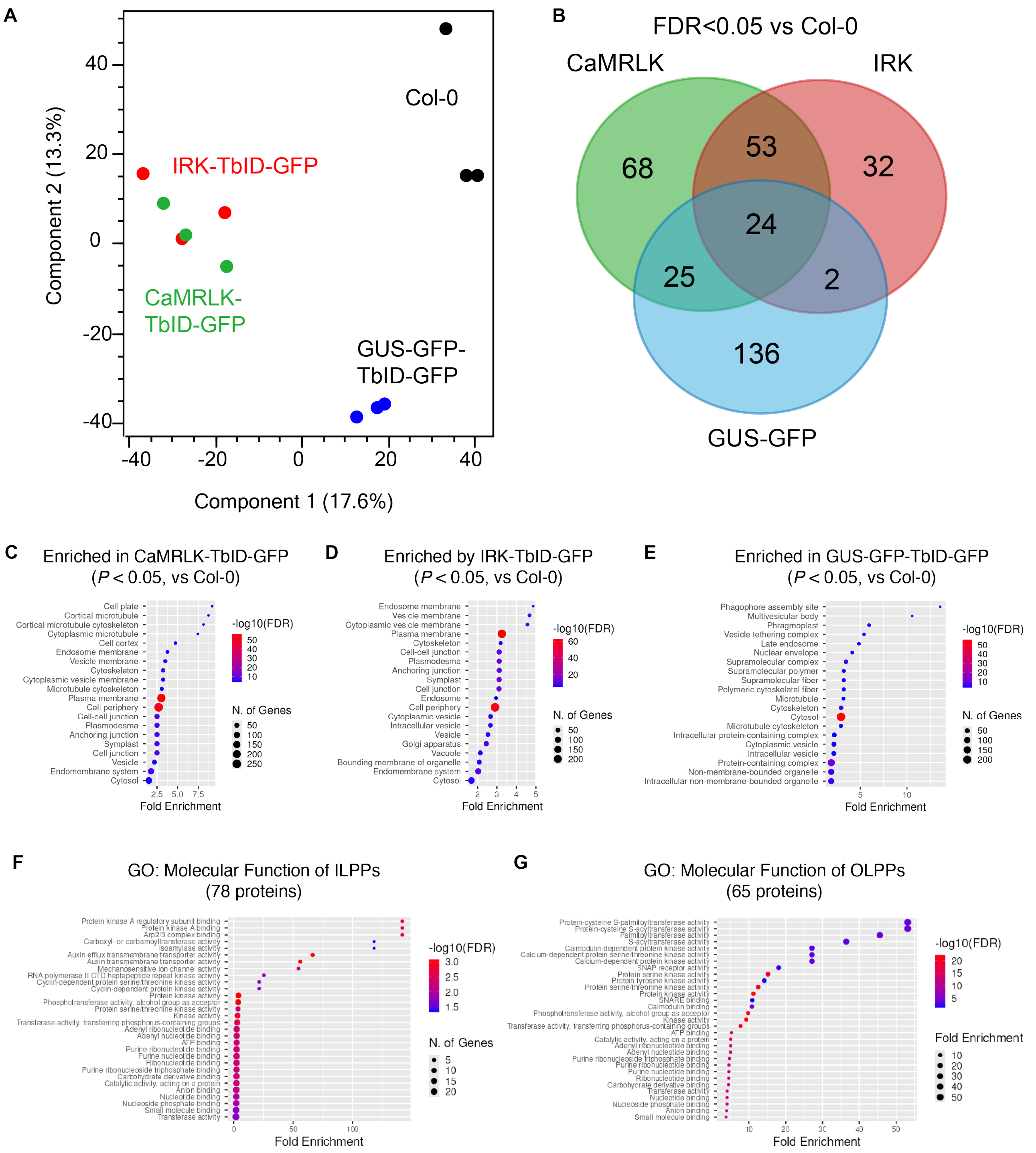


**Figure S2. Statistical analyses of the polarity-guided proximomes.**

(A) Principal component analysis (PCA) of the TbID-based polarity-guided proxitomes.

(B) Venn diagram showing the overlap among proteins significantly enriched in each proxitome relative to wild-type Col-0 (FDR < 0.05).

(C–E) Gene ontology (GO) enrichment analysis for the Cellular Component category of proteins significantly enriched by CaMRLK-TbID-GFP, IRK-TbID-GFP, or GUS-GFP-TbID relative to wild-type Col-0 (P < 0.05, Welch's t-test). GO enrichment analysis was performed using ShinyGO v0.741 (Ge et al. 2020)(http://bioinformatics.sdstate.edu/go74/).

(F and G) GO enrichment analysis for the Molecular Function category of ILPPs and OLPPs.

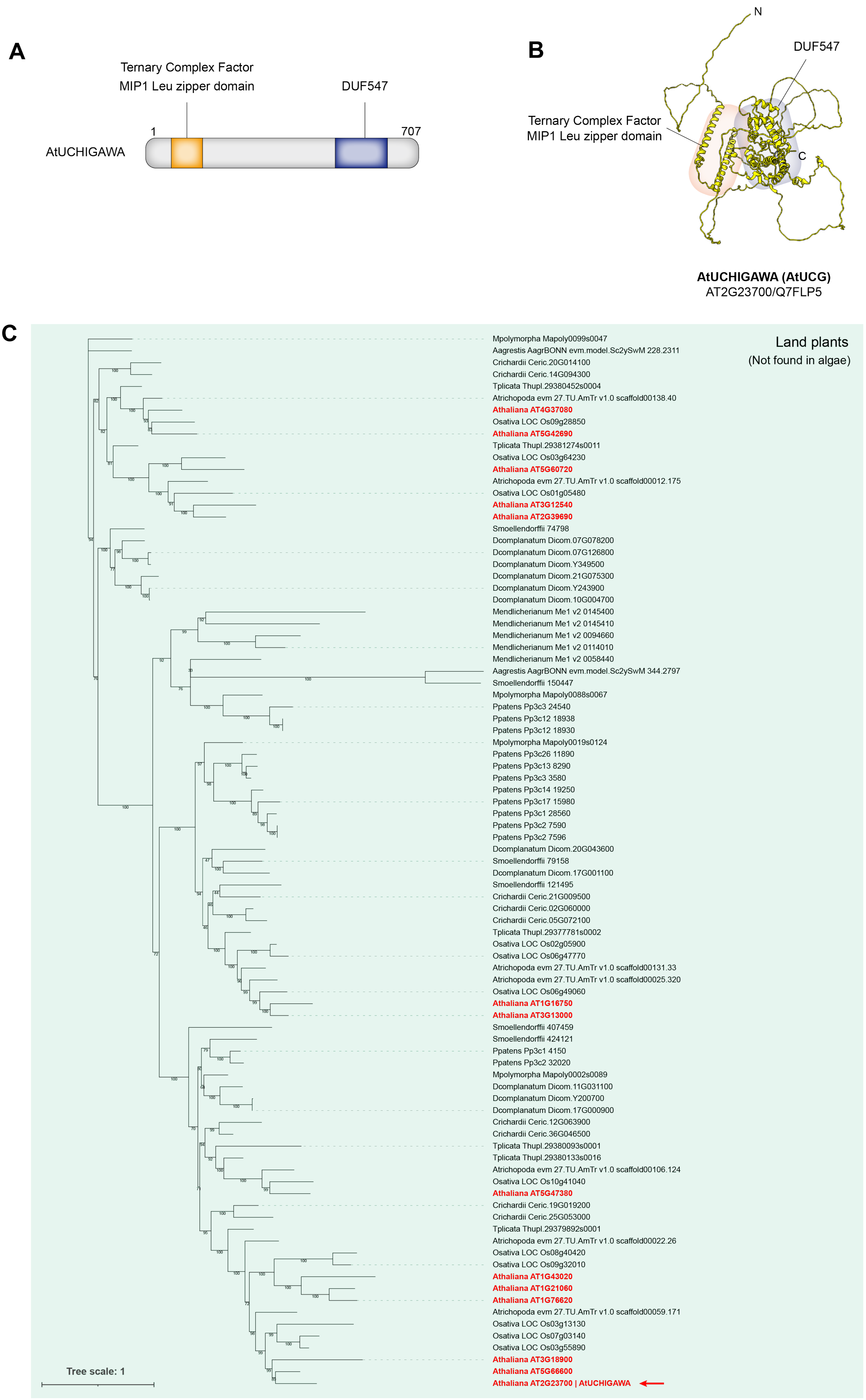
A
Ternary Complex Factor
MIP1 Leu zipper domain
DUF547
1
707
AtUCHIGAWA
B
N
DUF547
Ternary Complex Factor
MIP1 Leu zipper domain
C
AtUCHIGAWA (AtUCG)
AT2G23700/Q7FLP5
C
Land plants
(Not found in algae)
Mpolymorpha Mapoly0099s0047
Aagrestis AagrBONN evm.model.Sc2ySwM 228.2311
Crichardii Ceric.20G014100
Crichardii Ceric.14G094300
Tplicata Thupl.29380452s0004
Atrichopoda evm 27.TU.AmTr v1.0 scaffold00138.40
Athaliana AT4G37080
Osativa LOC Os09g28850
Athaliana AT5G42690
Tplicata Thupl.29381274s0011
Osativa LOC Os03g64230
Athaliana AT5G60720
Atrichopoda evm 27.TU.AmTr v1.0 scaffold00012.175
Osativa LOC Os01g05480
Athaliana AT3G12540
Athaliana AT2G39690
Smoellendorffii 74798
Dcomplanatum Dicom.07G078200
Dcomplanatum Dicom.07G126800
Dcomplanatum Dicom.Y349500
Dcomplanatum Dicom.21G075300
Dcomplanatum Dicom.Y243900
Dcomplanatum Dicom.10G004700
Mendlicherianum Me1 v2 0145400
Mendlicherianum Me1 v2 0145410
Mendlicherianum Me1 v2 0094660
Mendlicherianum Me1 v2 0114010
Mendlicherianum Me1 v2 0058440
Aagrestis AagrBONN evm.model.Sc2ySwM 344.2797
Smoellendorffii 150447
Mpolymorpha Mapoly0088s0067
Ppatens Pp3c3 24540
Ppatens Pp3c12 18938
Ppatens Pp3c12 18930
Mpolymorpha Mapoly0019s0124
Ppatens Pp3c26 11890
Ppatens Pp3c13 8290
Ppatens Pp3c3 3580
Ppatens Pp3c14 19250
Ppatens Pp3c17 15980
Ppatens Pp3c1 28560
Ppatens Pp3c2 7590
Ppatens Pp3c2 7596
Dcomplanatum Dicom.20G043600
Smoellendorffii 79158
Dcomplanatum Dicom.17G001100
Smoellendorffii 121495
Crichardii Ceric.21G009500
Crichardii Ceric.02G060000
Crichardii Ceric.05G072100
Tplicata Thupl.29377781s0002
Osativa LOC Os02g05900
Osativa LOC Os06g47770
Atrichopoda evm 27.TU.AmTr v1.0 scaffold00131.33
Atrichopoda evm 27.TU.AmTr v1.0 scaffold00025.320
Osativa LOC Os06g49060
Athaliana AT1G16750
Athaliana AT3G13000
Smoellendorffii 407459
Smoellendorffii 424121
Ppatens Pp3c1 4150
Ppatens Pp3c2 32020
Mpolymorpha Mapoly0002s0089
Dcomplanatum Dicom.11G031100
Dcomplanatum Dicom.Y200700
Dcomplanatum Dicom.17G000900
Crichardii Ceric.12G063900
Crichardii Ceric.36G046500
Tplicata Thupl.29380093s0001
Tplicata Thupl.29380133s0016
Atrichopoda evm 27.TU.AmTr v1.0 scaffold00106.124
Osativa LOC Os10g41040
Athaliana AT5G47380
Crichardii Ceric.19G019200
Crichardii Ceric.25G053000
Tplicata Thupl.29379892s0001
Atrichopoda evm 27.TU.AmTr v1.0 scaffold00022.26
Osativa LOC Os08g40420
Osativa LOC Os09g32010
Athaliana AT1G43020
Athaliana AT1G21060
Athaliana AT1G76620
Atrichopoda evm 27.TU.AmTr v1.0 scaffold00059.171
Osativa LOC Os03g13130
Osativa LOC Os07g03140
Osativa LOC Os03g55890
Athaliana AT3G18900
Athaliana AT5G66600
Athaliana AT2G23700 | AtUCHIGAWA
Tree scale: 1

**Figure S3. Phylogenetic tree and predicted three-dimensional structure of UCHIGAWA (UCG) protein.**

(A) Phylogenetic tree of UCHIGAWA (UCG) and its structurally related proteins identified by Foldseek (van Kempen et al. 2024)(https://search.foldseek.com/). Using the three-dimensional structure of AtUCG predicted by AlphaFold2, from *Arabidopsis thaliana*, 14 proteins were identified as structurally related proteins (*E*-value < $1 \times 10^{-20}$).

(B) Three-dimensional structure of AtUCG predicted by AlphaFold2. Approximate positions of the ternary complex factor MIP1 leucine zipper domain (PF14389) and DUF547 (PF04784) predicted by InterPro (https://www.ebi.ac.uk/interpro/) are highlighted. N, N-terminus; C, C-terminus.

(C) The phylogenetic tree was constructed using protein sequences homologous to AtUCG (AT2G23700) identified from representative green plant species. The query protein, AtUCG, is indicated by a red arrow. Protein sequences were aligned and analyzed using IQ-TREE3 with ultrafast bootstrap analysis (1,000 replicates). Bootstrap values are indicated at nodes. The tree was visualized using iTOL.

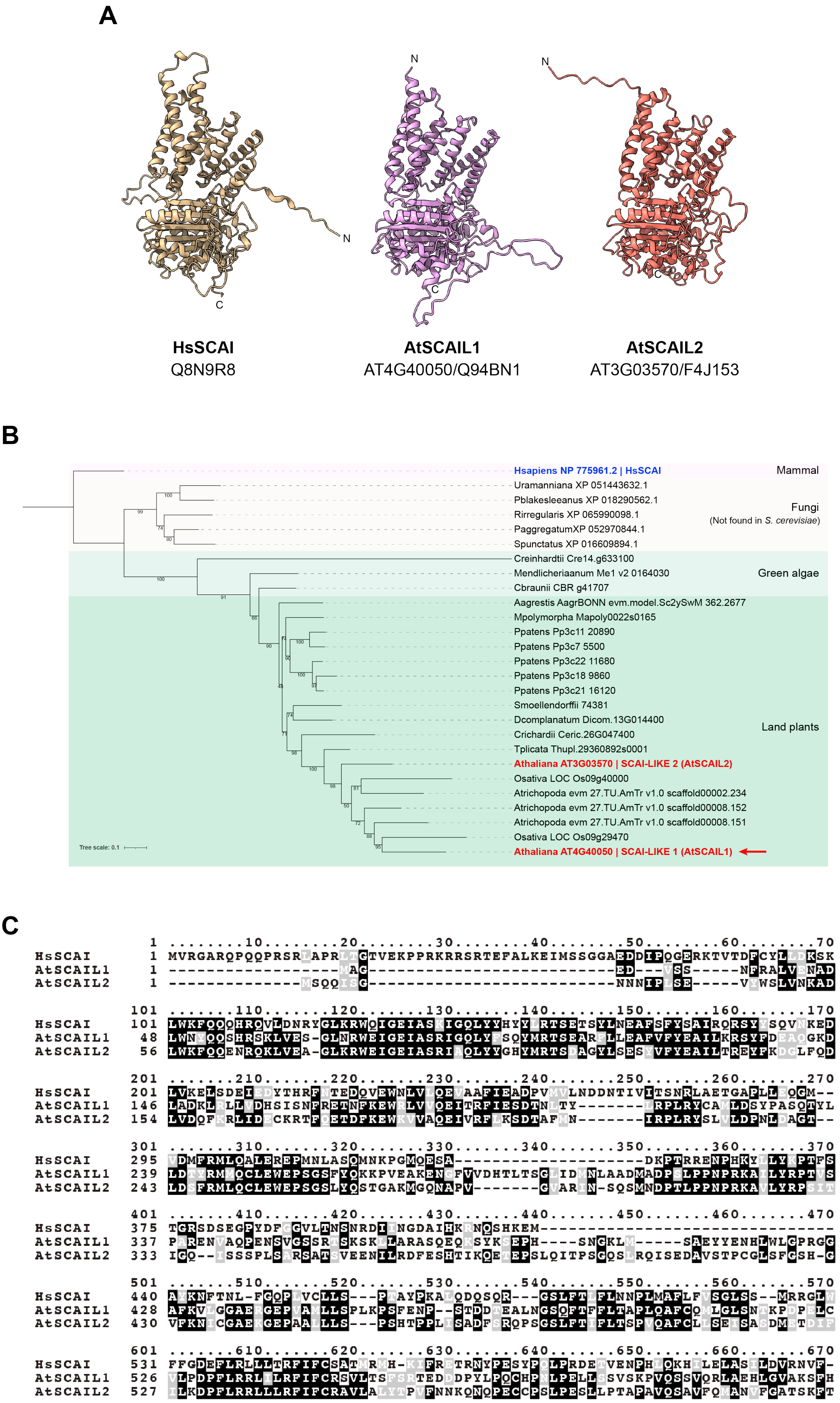
A
N
N
N
C
C
C
HsSCAI
Q8N9R8
AtSCAIL1
AT4G40050/Q94BN1
AtSCAIL2
AT3G03570/F4J153
B
Hsapiens NP 775961.2 | HsSCAI
Mammal
Uramanniana XP 051443632.1
Pblakesleeanus XP 018290562.1
Rirregularis XP 065990098.1
PaggregatumXP 052970844.1
Spunctatus XP 016609894.1
Fungi
(Not found in S. cerevisiae)
Creinhardtii Cre14.g633100
Mendlicheriaanum Me1 v2 0164030
Cbraunii CBR g41707
Green algae
Aagrestis AagrBONN evm.model.Sc2ySwM 362.2677
Mpolymorpha Mapoly0022s0165
Ppatens Pp3c11 20890
Ppatens Pp3c7 5500
Ppatens Pp3c22 11680
Ppatens Pp3c18 9860
Ppatens Pp3c21 16120
Smoellendorffii 74381
Dcomplanatum Dicom.13G014400
Crichardii Ceric.26G047400
Tplicata Thupl.29360892s0001
Athaliana AT3G03570 | SCAI-LIKE 2 (AtSCAIL2)
Osativa LOC Os09g40000
Atrichopoda evm 27.TU.AmTr v1.0 scaffold00002.234
Atrichopoda evm 27.TU.AmTr v1.0 scaffold00008.152
Atrichopoda evm 27.TU.AmTr v1.0 scaffold00008.151
Osativa LOC Os09g29470
Athaliana AT4G40050 | SCAI-LIKE 1 (AtSCAIL1)
Land plants
Tree scale: 0.1
C
1 ........10........20........30........40........50........60........70
HsSCAI 1 MVRGARQPQQPRSRLAPRLTGTVEKPPRKRRSRTEFALKEIMSSGGAEDDIPQGERKTVTDFCYLLDKSK
AtSCAIL1 1 ------------------MAG---------------------------ED---VSS-----NFRALVENAD
AtSCAIL2 1 ---------------MSQQISG---------------------------NNNIPLSE-----VIWSLVNKAD
101 .......110.......120.......130.......140.......150.......160.......170
HsSCAI 101 LWKFQQQHRQVLDNRYGLKRWQIGEIASKIGQLYYHYYLRTSETSYLNEAFSFYSAIRQRSYYSQVNKED
AtSCAIL1 48 LWNYQQSHRSKLVES-GLNRWEIGEIASRIGQLYFSQYMRTSEARFLLEAFVFYEAILKRSYFDEAQGKD
AtSCAIL2 56 LWKFQQENRQKLVEA-GLKRWEIGEIASRIAQLYYGHYMRTSDAGYLSESYVFYEAILTREYFKDGLFQD
201 .......210.......220.......230.......240.......250.......260.......270
HsSCAI 201 LVKELSDEIEDYTHRFLTEDQVEWNLVLQEVAAFIEADPVMVLNDDNTIVITSNRLAETGAPLLEQGM--
AtSCAIL1 146 LADKLRLLVDHSISNFRETNFKEWRLVVQEITRFIESDTNLTY-------IRPLRYCAMLDSYPASQTYL
AtSCAIL2 154 LVDQFKRLIDECKRTFSETDFKEWKVVAQEIVRFLKSDTAFMN--------IRPLRYSLVLDPNLPAGT--
301 .......310.......320.......330.......340.......350.......360.......370
HsSCAI 295 VDMFRMLQALEREPMNLASQMNKPGMQESA--------------------DKPTRRENPHKYLLYKPTFS
AtSCAIL1 239 LDTYRMMQCLEWEPSGSFYQKKPVEAKENSFVVDHTLTSGLIDMNLAADMADPSLPPNPRKAILYRPTVS
AtSCAIL2 243 LDSFRMLQCLEWEPSGSLYQSTGAKMGQNAPV-------GVARIN-SQSMNDPTLPPNPRKAVLYRPSIT
401 .......410.......420.......430.......440.......450.......460.......470
HsSCAI 375 TGRSDSEGPYDFCGVLTNSNRDIINGDAIHKRNQSHKEM-----------------------------
AtSCAIL1 337 PARENVAQPENSVGSSRTSKSKLLARASQEQKSYKSEPH----SNGKLM-----SAEYYENHLWLGPRGG
AtSCAIL2 333 IGQ--ISSSPLSARSATSVEENILRDFESHTIKQETEPSLQITPSGQSLRQISEDAVSTPCGLSFGSH-G
501 .......510.......520.......530.......540.......550.......560.......570
HsSCAI 440 ALKNFTNL-FGQPLVCLLS---PTAYPKALQDQSQR---GSLFTLFLNNPLMAFLFVSGLSS--MRRGLW
AtSCAIL1 428 AFKVLGGAERGEPVALLSPLKPSFENP--STDDTEALNGSQFTFFLTAPLQAFCQMLGLSNTKPDPELC
AtSCAIL2 430 VFKNICGAEKGEPAALLS---PSHTPPLISADFSRQPSGSLFTIFLTSPVQAFCLLSEISASDMLTDIF
601 .......610.......620.......630.......640.......650.......660.......670
HsSCAI 531 FFGDEFLRLLLTRFIFCSATMRMH-KIFRETRNYPESYPQLPRDETVENPHLQKHILELASHLDVRNVF-
AtSCAIL1 526 VLPDPFLRRLILRFIFCRSVLTSFSRTEDDDPYLPQCHPNLPELLSSVSKPVQSSVQRLAEHLGVAKSFH
AtSCAIL2 527 ILKDPFLRRLLLRFIFCRAVLALYTPVFNNKQNQPECCPSLPESLLPTAPAVQSAVFQMANVFCATSKFT

**Figure S4. Predicted three-dimensional structure, phylogenetic tree, and multiple sequence alignment of human SCAI and Arabidopsis SCAI-LIKE (SCAIL) proteins.**

(A) Three-dimensional structure of *Homo sapiens* SCAI, Arabidopsis SCAI-LIKE 1 (SCAIL1), Arabidopsis SCAIL2 predicted by AlphaFold2. N, N-terminus; C, C-terminus.

(B) Phylogenetic tree of SCAI and SCAIL proteins in angiosperms (*Arabidopsis thaliana*, *Oryza sativa*, and *Amborella trichopoda*), a gymnosperm (*Thuja plicata*), a fern (*Ceratopteris richardii*), Lycophyte (*Selaginella moellendorffii*), a moss (*Physcomitrella patens*), a liverwort (*Marchantia polymorpha*), a charophyte alga (*Chara vulgaris*), fungi (*Spizellomyces punctatus*, *Polychytrium aggregatum*, *Phycomyces blakesleeanus*, *Umbelopsis ramanniana*, and *Rhizophagus irregularis)*, and human (*Homo sapiens*). SCAI-LIKE proteins were not identified in *Saccharomyces cerevisiae.* The phylogenetic tree was constructed using protein sequences homologous to AtSCAIL1 (AT4G40050) identified from representative eukaryotic lineages. The query protein, AtSCAIL1, is indicated by a red arrow. Protein sequences were aligned and analyzed using IQ-TREE3 with ultrafast bootstrap analysis (1,000 replicates). Bootstrap values are indicated at nodes. The tree was visualized using iTOL.

(C) Multiple sequence alignment of HsSCAI, AtSCAIL1, and AtSCAIL2, calculated by the MUSCLE algorithm using Galaxy (ver. 0.0.11; https://usegalaxy.eu/).

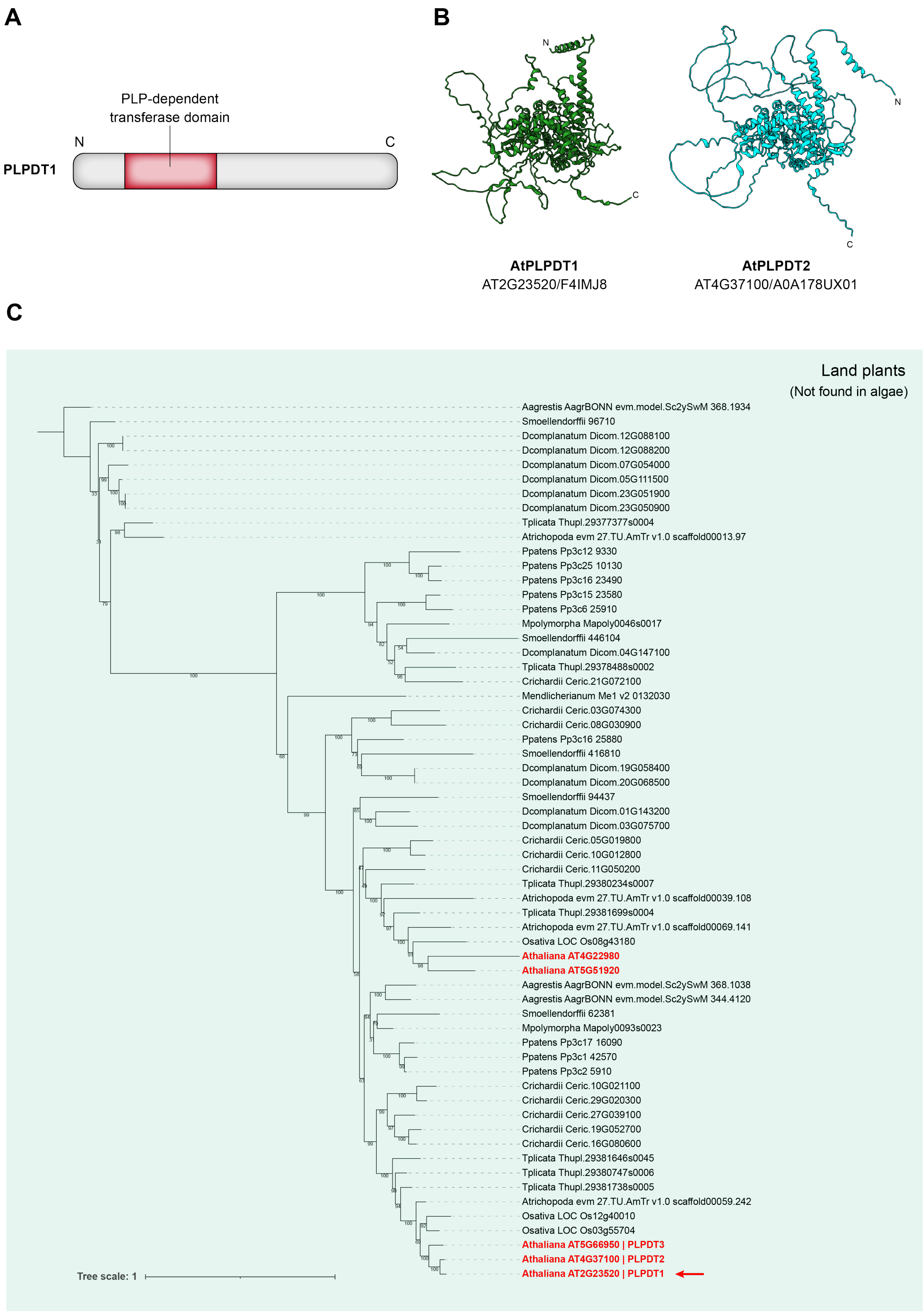
A
PLP-dependent transferase domain
N
C
PLPDT1
B
N
C
N
C
AtPLPDT1
AT2G23520/F4IMJ8
AtPLPDT2
AT4G37100/A0A178UX01
C
Land plants
(Not found in algae)
Aagrestis AagrBONN evm.model.Sc2ySwM 368.1934
Smoellendorffii 96710
Dcomplanatum Dicom.12G088100
Dcomplanatum Dicom.12G088200
Dcomplanatum Dicom.07G054000
Dcomplanatum Dicom.05G111500
Dcomplanatum Dicom.23G051900
Dcomplanatum Dicom.23G050900
Tplicata Thupl.29377377s0004
Atrichopoda evm 27.TU.AmTr v1.0 scaffold00013.97
Ppatens Pp3c12 9330
Ppatens Pp3c25 10130
Ppatens Pp3c16 23490
Ppatens Pp3c15 23580
Ppatens Pp3c6 25910
Mpolymorpha Mapoly0046s0017
Smoellendorffii 446104
Dcomplanatum Dicom.04G147100
Tplicata Thupl.29378488s0002
Crichardii Ceric.21G072100
Mendlicherianum Me1 v2 0132030
Crichardii Ceric.03G074300
Crichardii Ceric.08G030900
Ppatens Pp3c16 25880
Smoellendorffii 416810
Dcomplanatum Dicom.19G058400
Dcomplanatum Dicom.20G068500
Smoellendorffii 94437
Dcomplanatum Dicom.01G143200
Dcomplanatum Dicom.03G075700
Crichardii Ceric.05G019800
Crichardii Ceric.10G012800
Crichardii Ceric.11G050200
Tplicata Thupl.29380234s0007
Atrichopoda evm 27.TU.AmTr v1.0 scaffold00039.108
Tplicata Thupl.29381699s0004
Atrichopoda evm 27.TU.AmTr v1.0 scaffold00069.141
Osativa LOC Os08g43180
Athaliana AT4G22980
Athaliana AT5G51920
Aagrestis AagrBONN evm.model.Sc2ySwM 368.1038
Aagrestis AagrBONN evm.model.Sc2ySwM 344.4120
Smoellendorffii 62381
Mpolymorpha Mapoly0093s0023
Ppatens Pp3c17 16090
Ppatens Pp3c1 42570
Ppatens Pp3c2 5910
Crichardii Ceric.10G021100
Crichardii Ceric.29G020300
Crichardii Ceric.27G039100
Crichardii Ceric.19G052700
Crichardii Ceric.16G080600
Tplicata Thupl.29381646s0045
Tplicata Thupl.29380747s0006
Tplicata Thupl.29381738s0005
Atrichopoda evm 27.TU.AmTr v1.0 scaffold00059.242
Osativa LOC Os12g40010
Osativa LOC Os03g55704
Athaliana AT5G66950 | PLPDT3
Athaliana AT4G37100 | PLPDT2
Athaliana AT2G23520 | PLPDT1
Tree scale: 1

**Figure S5. Predicted three-dimensional structure, schematic illustration of protein domain architecture, and phylogenetic tree of PLPDT1/2.**

(A) Three-dimensional structure of PLPDT1 and PLPDT2 predicted by AlphaFold2. N, N-terminus; C, C-terminus.

(B) Location of the characteristic domain for PLP-dependent transferases (SSF53383) within PLPDT1 predicted by InterPro.

(C) Phylogenetic tree of PLPDT proteins in angiosperms (*Arabidopsis thaliana*, *Oryza sativa*, and *Amborella trichopoda*), a gymnosperm (*Thuja plicata*), a fern (*Ceratopteris richardii*), Lycophyte (*Selaginella moellendorffii*), a moss (*Physcomitrella patens*), a liverwort (*Marchantia polymorpha*), a charophyte alga (*Chara vulgaris*). The phylogenetic tree was constructed using protein sequences homologous to AtPLPDT (AT2G23520) identified from representative eukaryotic lineages. The query protein, AtPLPDT1, is indicated by a red arrow. Protein sequences were aligned and analyzed using IQ-TREE3 with ultrafast bootstrap analysis (1,000 replicates). Bootstrap values are indicated at nodes. The tree was visualized using iTOL.

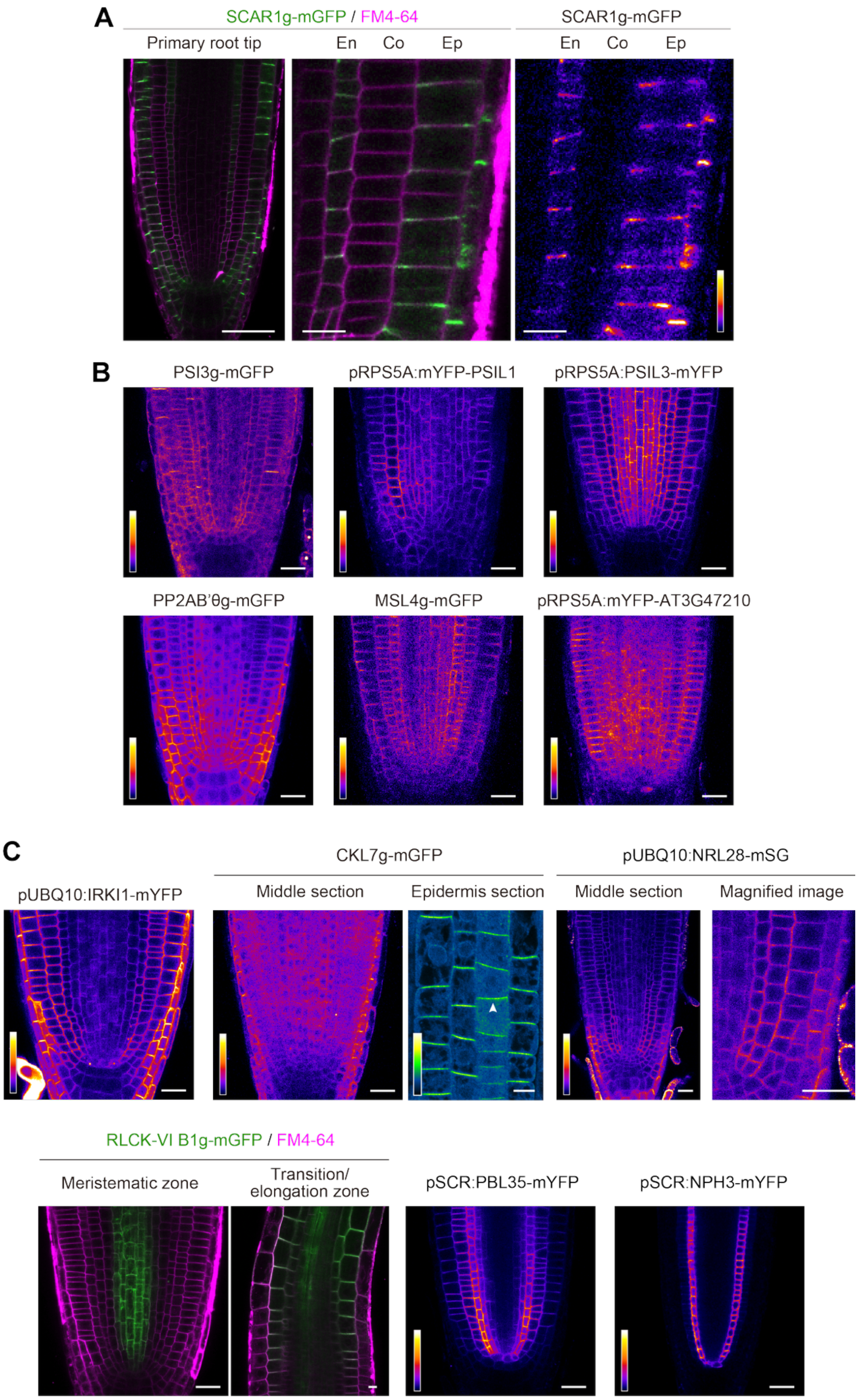


**Figure S6. ILPP and OLPP candidates which exhibited a bipolar or apolar subcellular localization *in planta*.**

(A) Confocal images of SCAR1/WAVE1-mGFP expressed under the control of native promoters. Four-day-old plants of $T_2$ generation were used for the imaging. Scale bar: 50 µm (left) and 10 µm (middle and right panels).
(B) Confocal images of the ILPP candidates including PSI3, PSIL1, PSIL3, PP2A B'θ subunit, MSL4, and AT3G47210. Proteins except AT3G47210 were fused with mGFP and expressed under the control of native promoters. AT3G47210 was fused with mYFP N-terminally and expressed under the control of the *RPS5A* promoter. Four-day-old plants of $T_2$ generation were used for the imaging. Scale bar: 20 µm.
(C) Confocal images of the OLPP candidates including CKL7, RLCK-VI B1, PBL35, and PHGAP1 fused with mGFP, NRL28 fused with StayGold$^{E138D}$ (mSG), and NPH3 fused with mYFP. CKL7, RLCK-VI B1, PBL35, and PHGAP1 fused with mGFP were expressed under the control of native promoters. NRL28-mSG and NPH3-mYFP were expressed under the control of *UBQ10* and *SCR* promoters, respectively. IRKI1 was fused with mYFP and ectopically expressed under the control of the *RPS5A* promoter. Except for NRL28-mSG, 4-day-old $T_2$-generation plants were used for the imaging. For the imaging of NRL28-mSG, $T_1$ generation of transgenic plants were grown on antibiotic-containing selection medium for 4 days, transferred to antibiotic-free MGRL medium, and grown for 2 days before imaging. Scale bar: 20 µm.

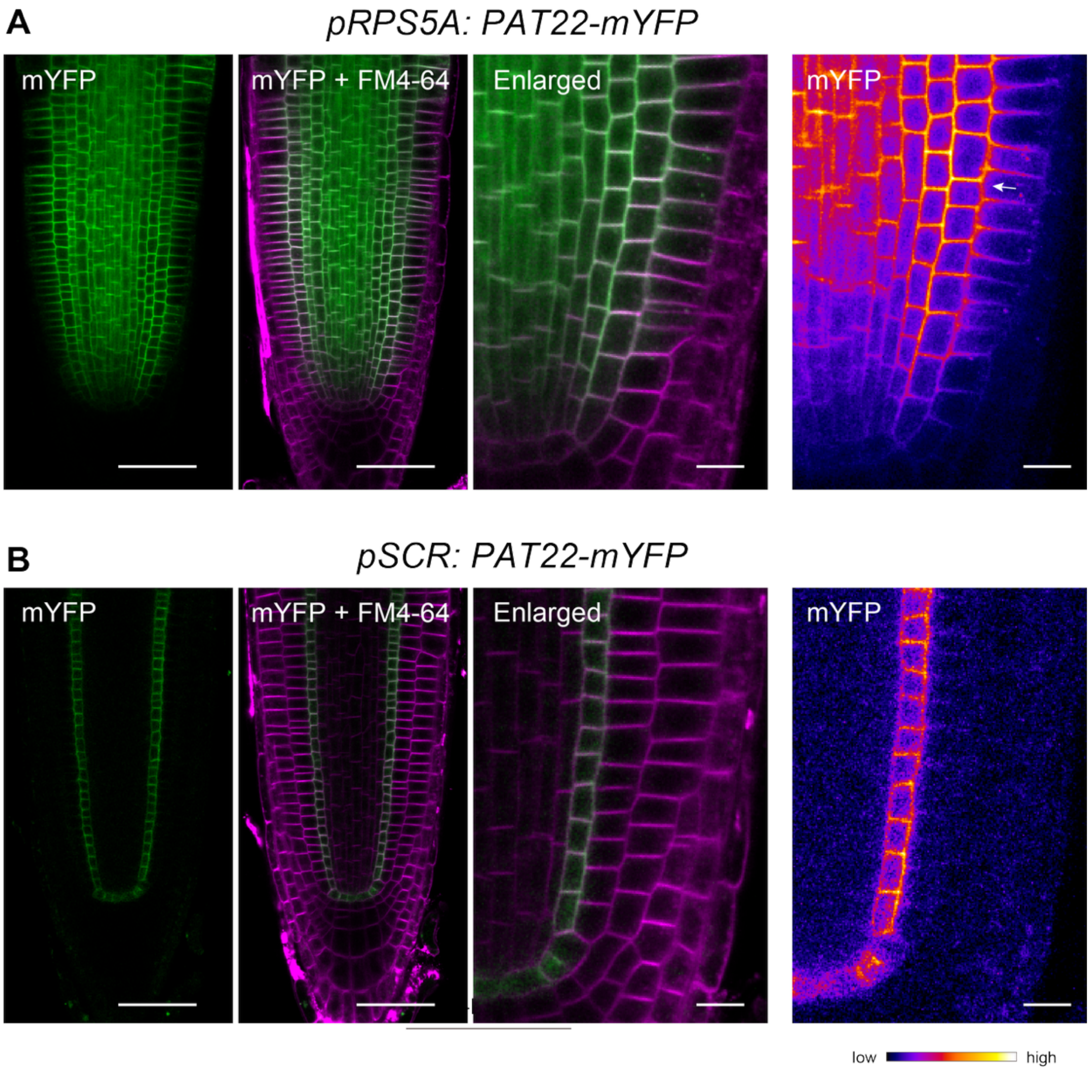


**Figure S7. PAT22 is weakly polarized in the epidermal cells.**

Confocal images of PAT22-mYFP expressed under the control of *RPS5A* (A) and *SCR* (B) promoters. The arrow indicates polarity orientation. Scale bars: 50 µm (left panels), 10 µm (right magnified panels).

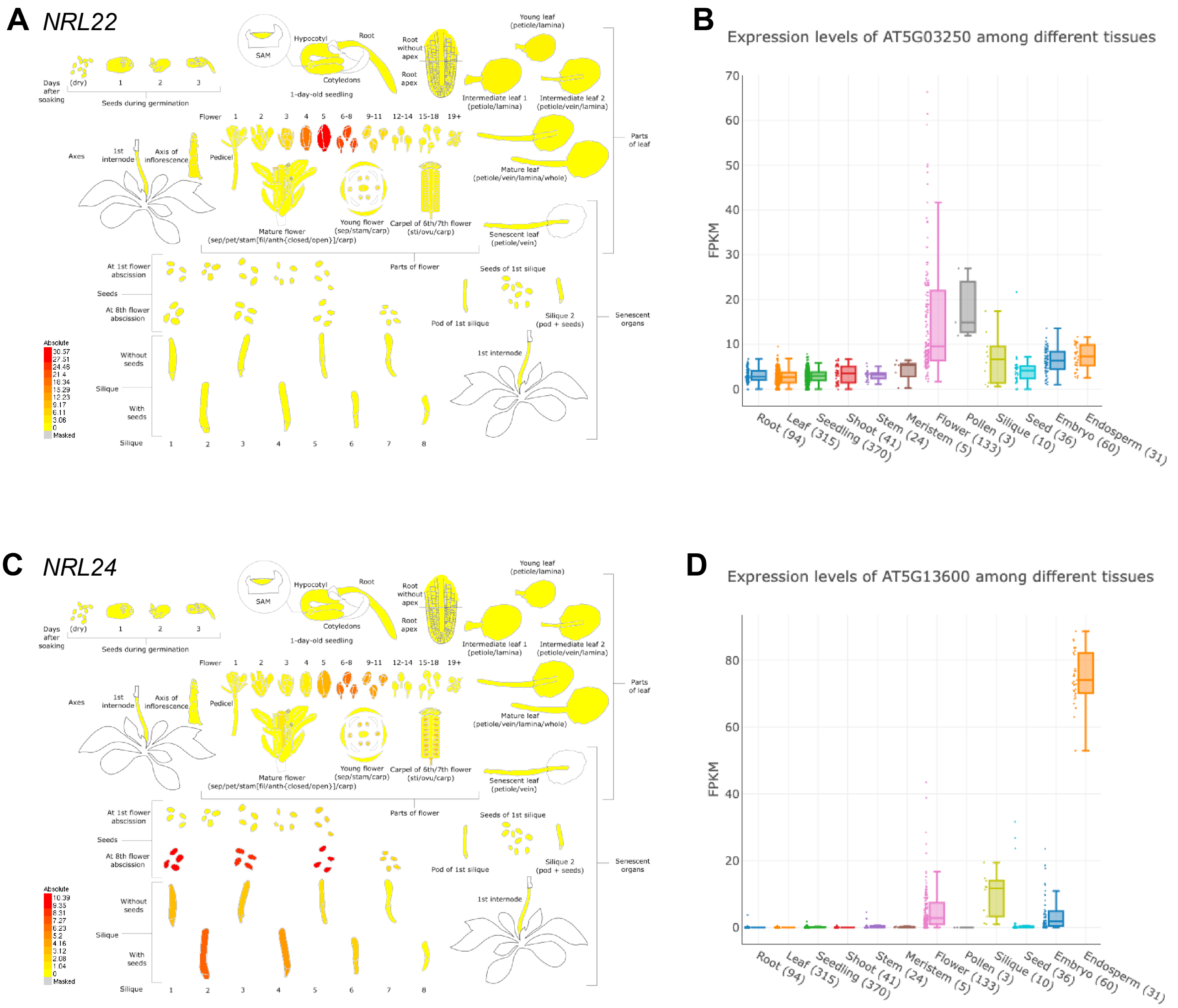


**Figure S8. Public transcriptome data indicate the preferential expression of *NRL22* and *NRL24* in reproductive organs in *Arabiodpsis thaliana.***

(A and C) Tissue-specific transcript abundance of *NRL22* (A) and *NRL24* (C) determined by a single cell RNA-seq (scRNA-seq), so-called Klepikova Atlas[37], obtained from TAIR (https://www.arabidopsis.org/).

(B and D) Normalized tissue-specific transcript level of *NRL22* (B) and *NRL24* (D) based on public RNA-seq libraries obtained from Arabidopsis RNA-seq Database (https://plantrnadb.com/athrdb/).

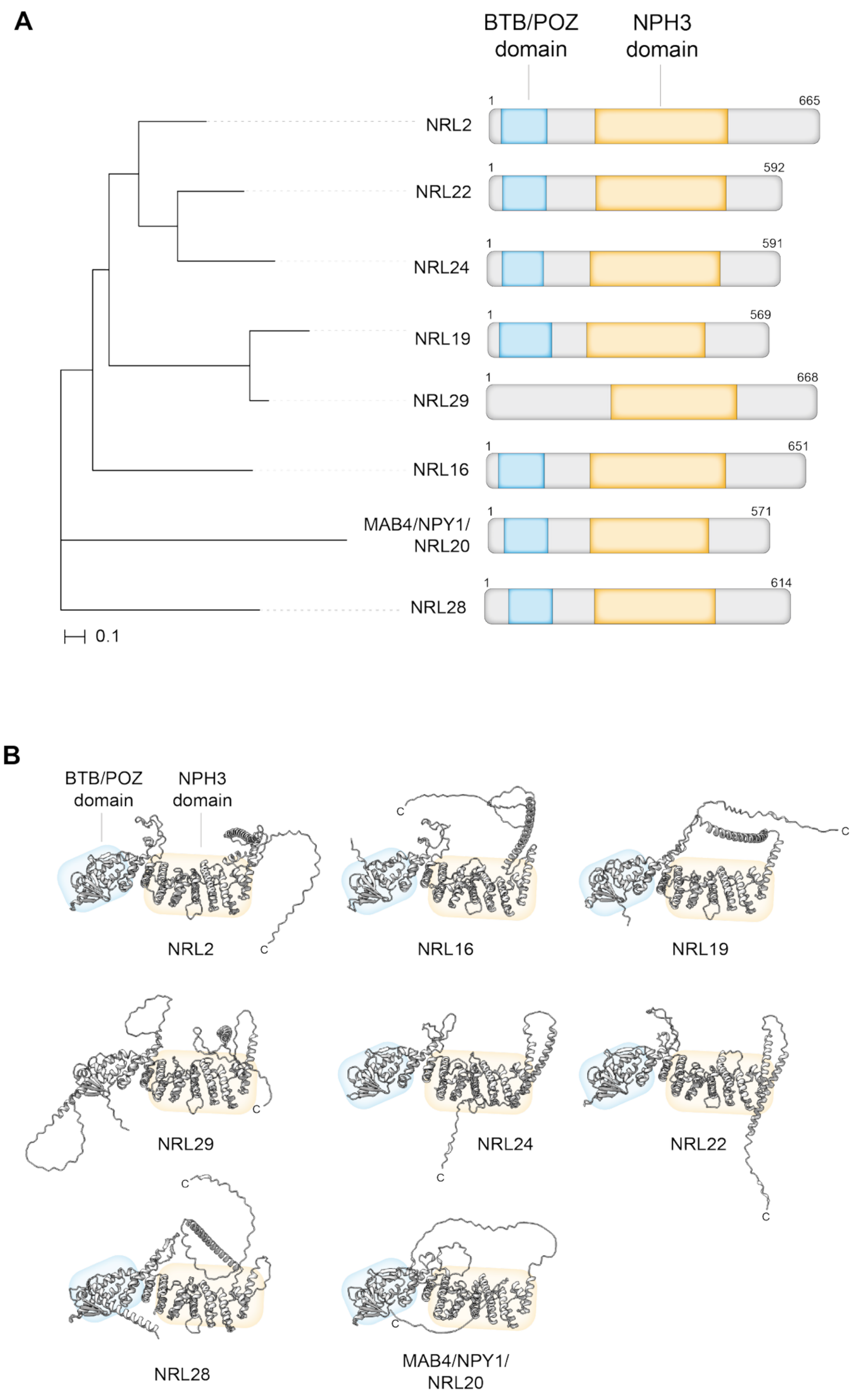


**Figure S9. NRLs are conserved in land plants.**

(A) Phylogenetic tree of NPT-clade NRLs with MAB4 and NRL28 (outgroups). Corresponding protein domain architectures are indicated on the right. BTB/POZ (PF00651) and NPH3 (PF03000) domains, predicted by InterPro, are indicated in blue and orange, respectively.

(B) Three-dimensional structures of NPT-clade NRLs with MAB4 and NRL28. Approximate positions of BTB/POZ and NPH3 domains are indicated. NRL29 has a longer N-terminus and BTB/POZ domain was not predicted by InterPro.

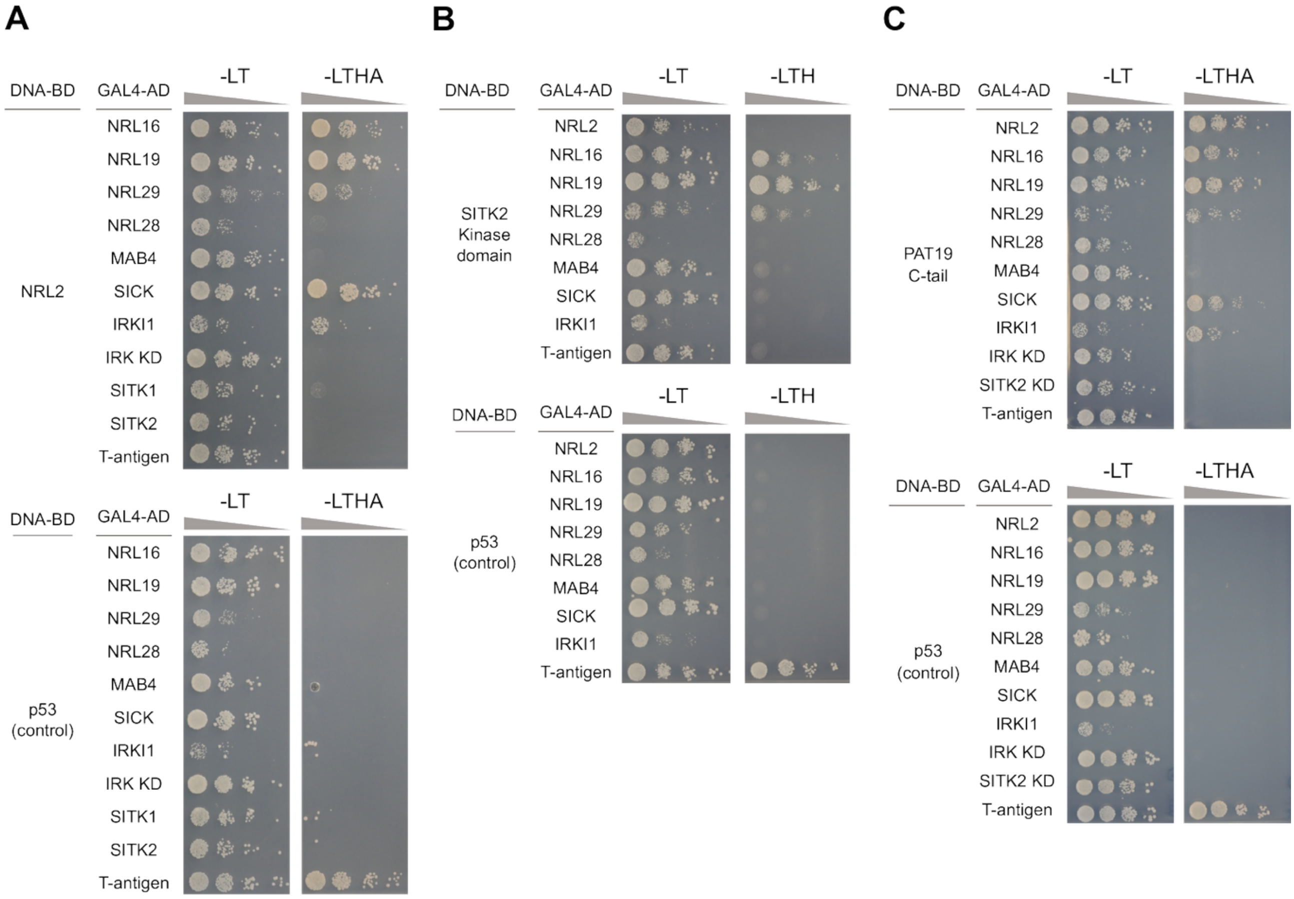


**Figure S10. Yeast two-hybrid assay of LPC components**

Yeast cultures were adjusted to the equivalent optical density ($OD_{600}$), serially diluted 10-fold (1, $10^{-1}$, $10^{-2}$, and $10^{-3}$), and spotted onto the SD/-Trp/-Leu (control), SD/-Trp/-Leu/-His (-LTH), or SD/-Trp/-Leu/-His/-Ade (-LTHA) medium. p53 and SV40 T-antigen were used as control of the bait and prey, respectively.

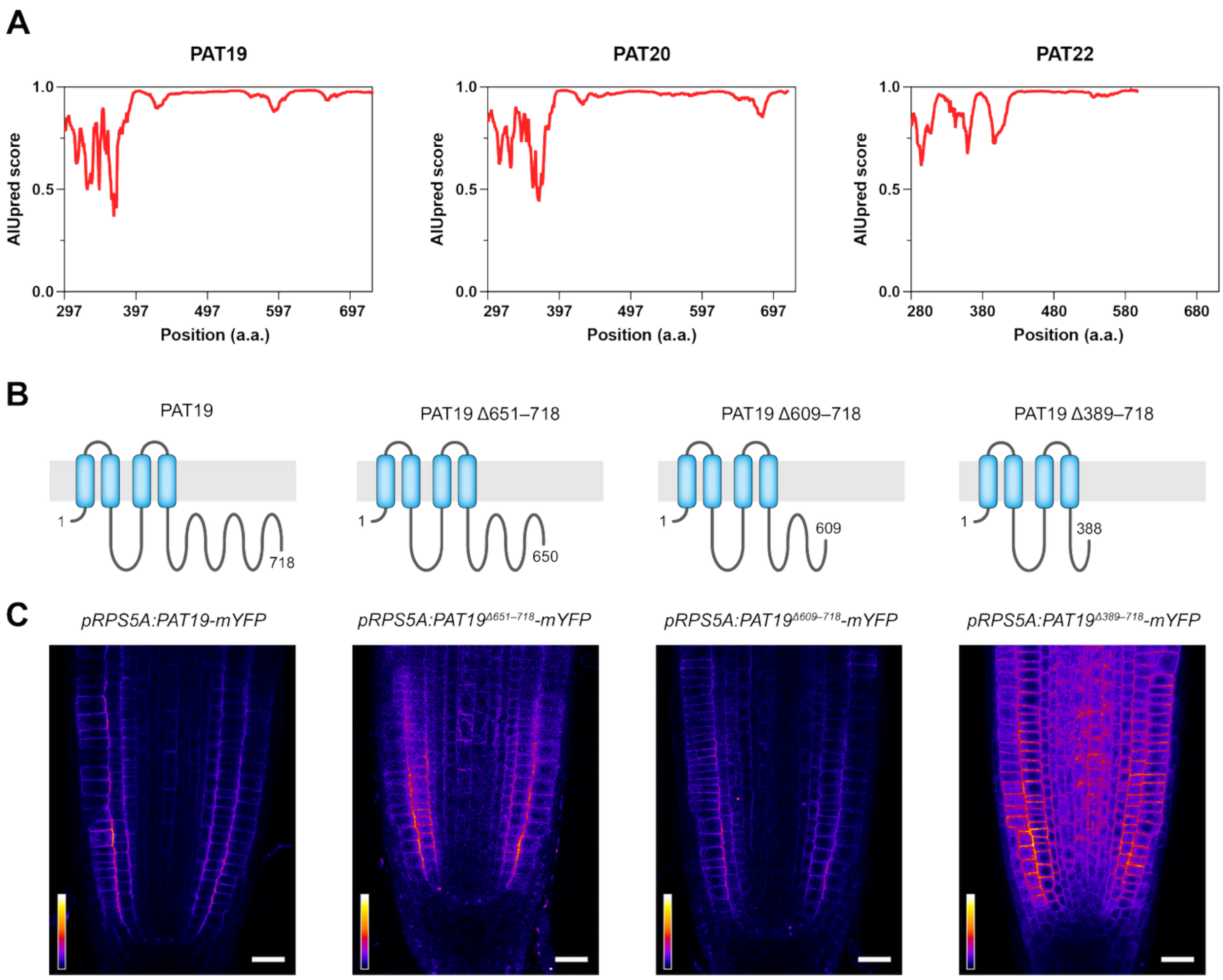


**Figure S11. Polar localization of PAT19 requires its C-terminal region.**

(A) Schematic illustration of putative intrinsically disordered regions (IDRs) within the C-terminal region of PAT19 predicted by AIUPred.

(B) Schematic illustration of truncation series of PAT19 tested in the present study.

(C) Confocal images of the PAT19 variants fused with mYFP which have the native or shorter C-terminal regions in 4-day-old Arabidopsis seedlings. Transgenes were expressed under the control of *RPS5A* promoter. Scale bar: 20 µm.

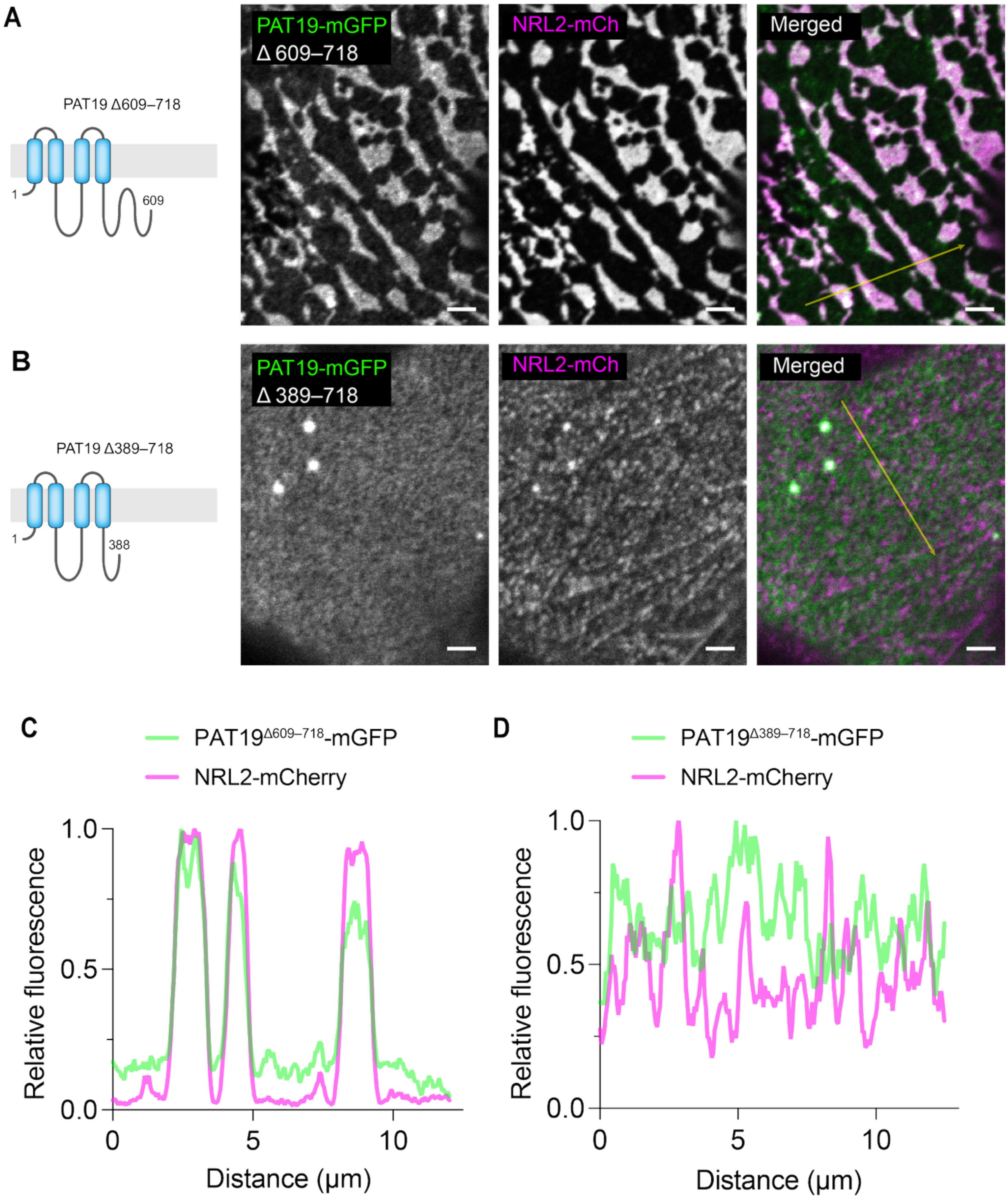


**Figure S12. Colocalization of PAT19 with NRL2 requires its C-terminal region.**

(A and B) Confocal images of truncated variants of PAT19-mGFP co-expressed with NRL2-mCherry at the outer surface of *N. benthamiana* epidermal cells. Scale bar: 20 μm.

(C and D) Fluorescence intensity along the arrows with 3-pixel width indicated in the confocal images in A and B.

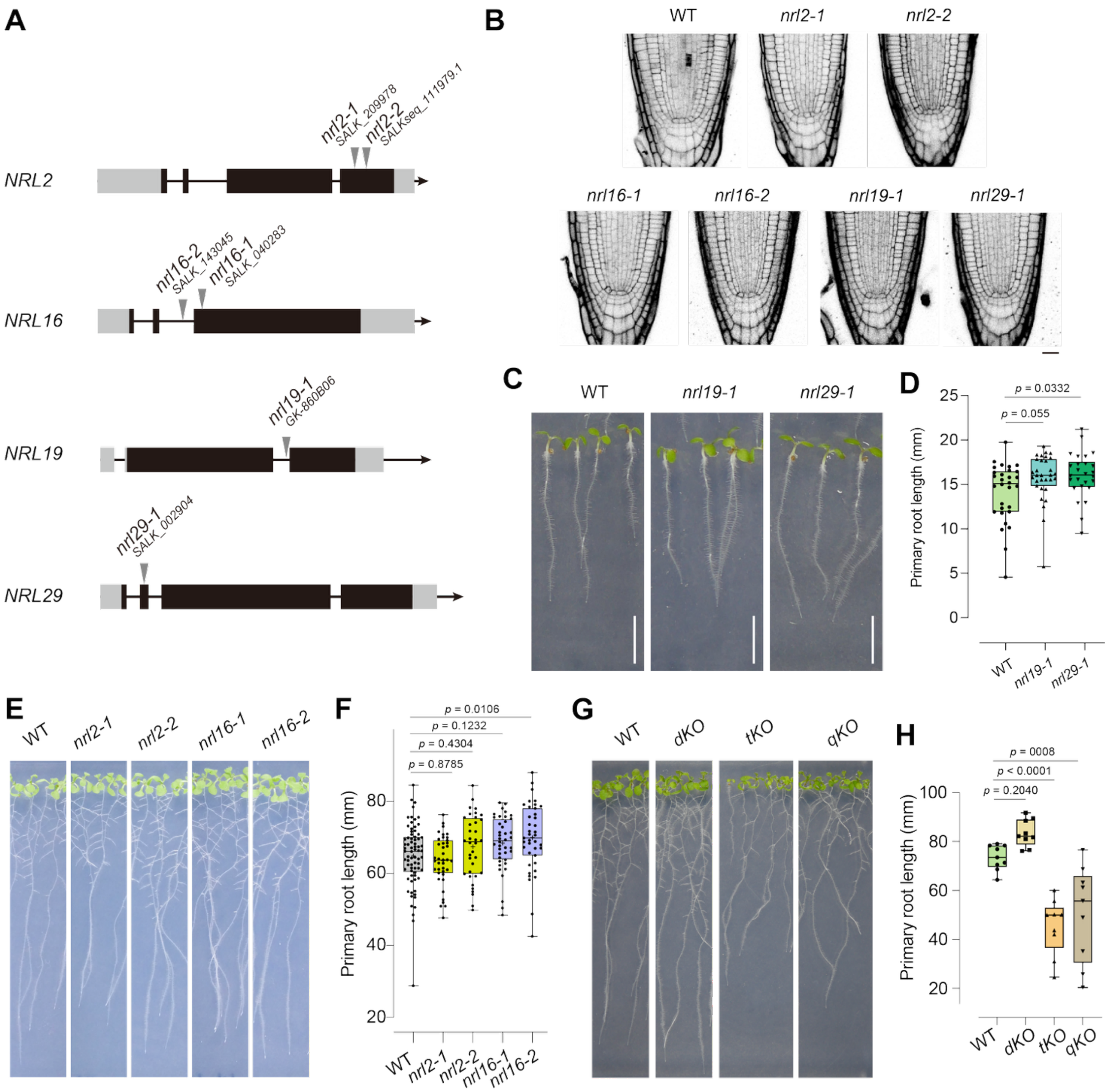


**Figure S13. T-DNA insertion lines of *NRL2*, *NRL16*, *NRL19*, and *NRL29*.**

(A) Gene architecture. Grey regions indicate 5'UTR and 3'UTR. Wide black bars indicate exons. Narrow bars indicate introns. Positions of T-DNA insertion are described by arrowheads.

(B) Confocal images of the root apical meristem of single T-DNA insertion mutants. Plants were grown vertically for 4 days on MGRL medium. Cell walls were stained with propidium iodide (PI). Scale bar: 10 µm.

(C) Growth phenotype of wild type, *nrl19-1* and *nrl29-1*. Plants were grown vertically for 5 days on MGRL medium. Scale bar: 5 mm.

(D) Quantification of primary root length of wild type, *nrl19-1*, and *nrl29-1* seedlings. *P* values were determined by Dunnett's multiple comparison test with WT.

(E) Growth phenotype of wild type, *nrl2-1*, *nrl2-2*, *nrl16-1*, and *nrl16-2* seedlings grown for 9 days on MGRL medium.

(F) Quantification of primary root length of wild type, *nrl2-1*, *nrl2-2*, *nrl16-1*, and *nrl16-2. P* values were determined by Dunnett's multiple comparison test with WT. *n* = 73 (WT), 39 (*nrl2-1*), 36 (*nrl2-2*), 40 (*nrl16-1*), and 39 (*nrl16-2*).
(G) Growth phenotype of wild type, *nrl2-2;nrl16-2* (dKO), *nrl2-2;nrl16-2;nrl29-1* (tKO), and *nrl2;nrl16;nrl19;nrl24;nrl29;NRL22*$^{\Delta 9aa}$ (qKO) seedlings grown for 9 days on MGRL medium.
(H) Quantification of primary root length of wild type, *nrl2-2;nrl16-2* (dKO), *nrl2-2;nrl16-2;nrl29-1* (tKO), and *nrl2;nrl16;nrl19;nrl24;nrl29;NRL22*$^{\Delta 9aa}$ (qKO) seedlings *P* values were determined by Dunnett's multiple comparison test with WT. *n* = 9 for each sample.

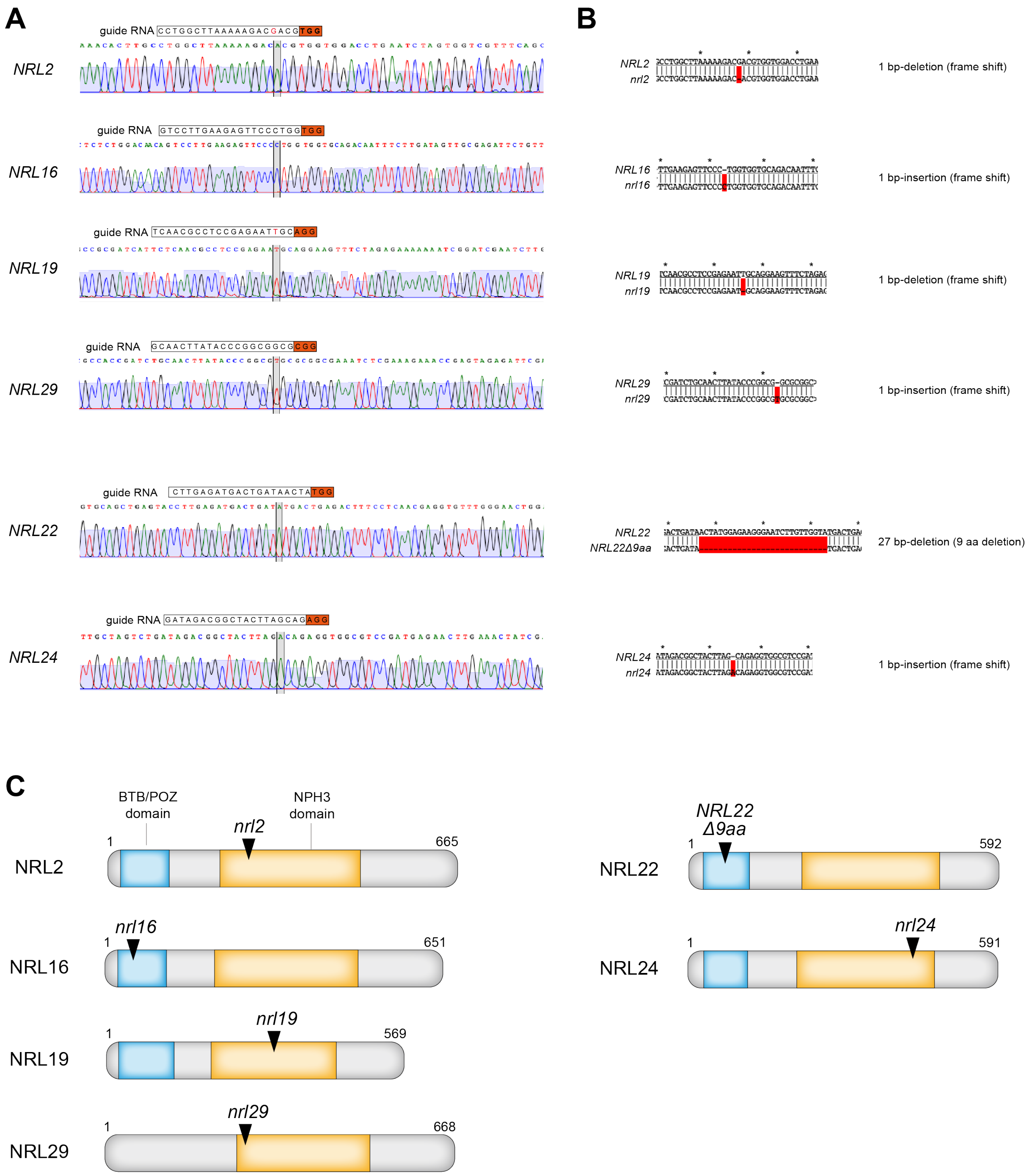


**Figure S14. Genome editing of NPT-clade *NRLs*.**

(A) Guide RNA and PAM sequence and chromatogram of Sanger sequencing.

(B) Mutation types. DNA sequence alignments of wild-type (top) and mutated (bottom) sequences. Mismatches/gaps are highlighted in red.

(C) Positions of the mutations within the protein domain architecture.

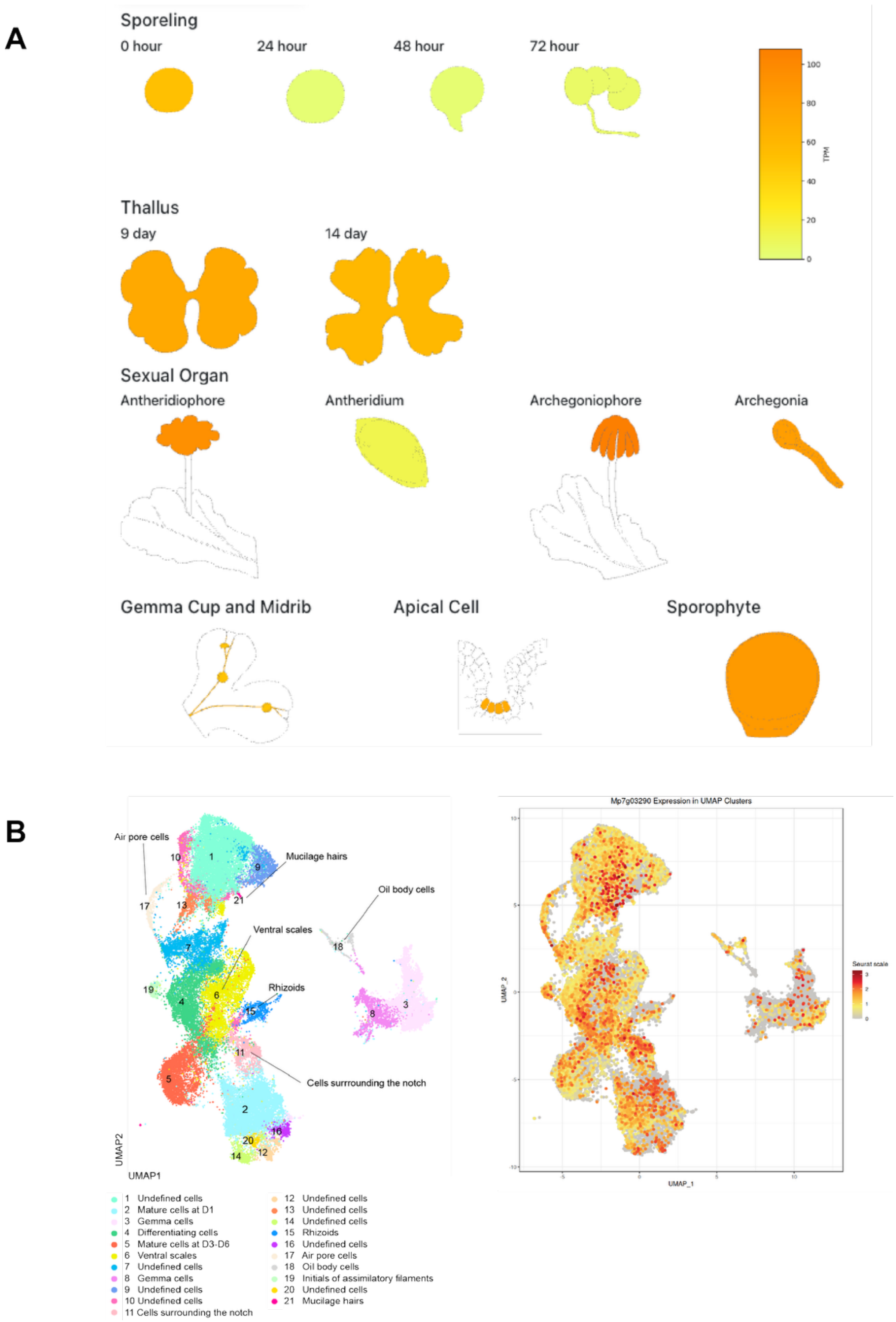


**Figure S15. Mp*NPT* is broadly expressed across the organs.**

(A) Tissue-specific RNA-seq data of Mp*NPT* gene obtained from MaprolBase (https://mbex.marchantia.info/).

(B) scRNA-seq data of Mp*NPT* gene [38].

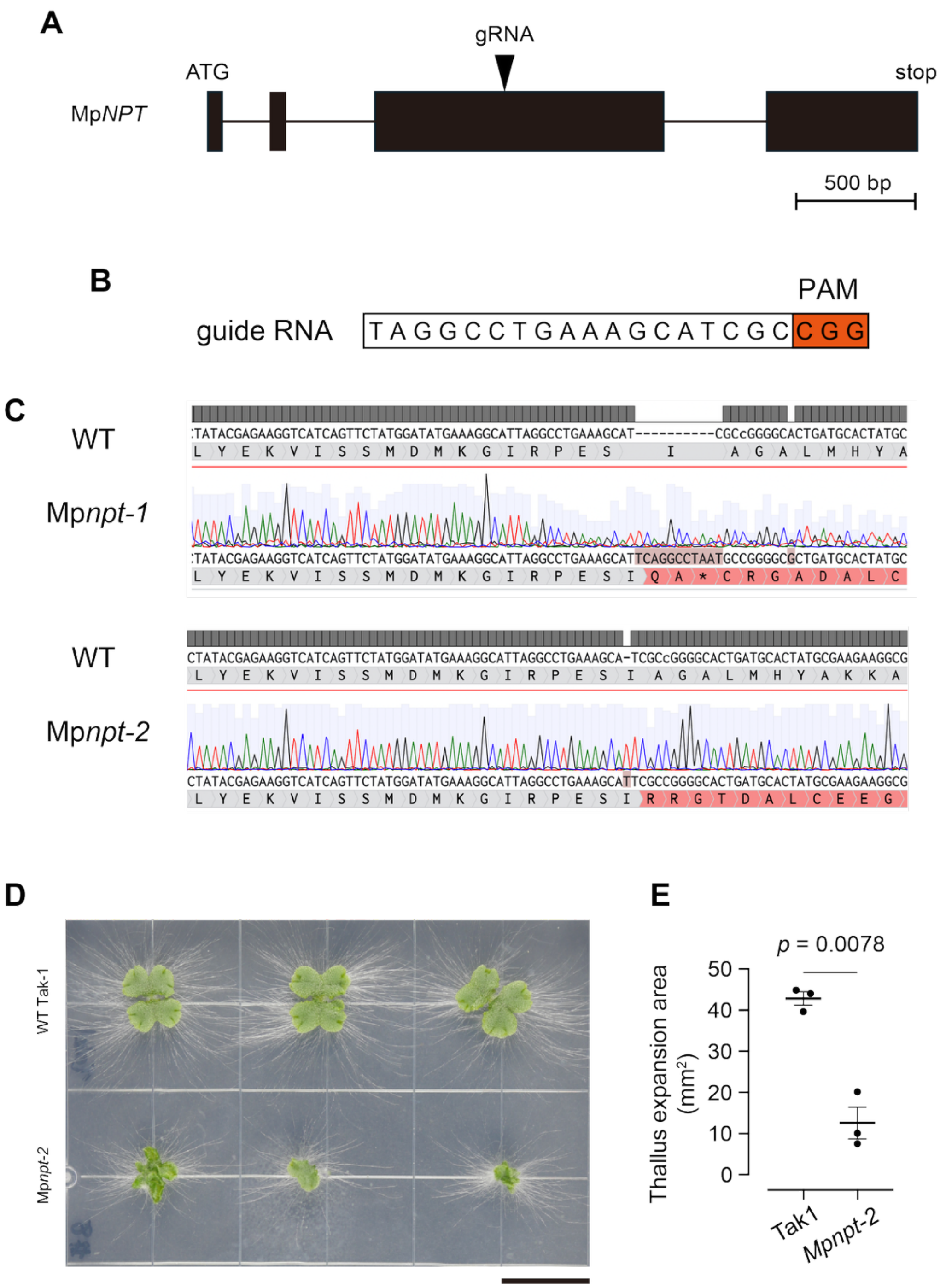


**Figure S16. Genome editing of Mp*NPT*.**

(A) Schematic representation of the Mp*NPT* gene and the positions of the sgRNA target sites. Boxes and lines indicate exons and introns, respectively. The orange arrowhead indicates the sgRNA target site.

(B) Nucleotide sequence surrounding the sgRNA target site. The sgRNA target sequence is shown in blue, and the protospacer-adjacent motif (PAM) is underlined in orange.

(C) Nucleotide sequences of the 11-bp and 1-bp insertions within Mp*NPT* gene and representative Sanger sequencing chromatograms of the genes amplified from Mp*npt-1* and Mp*npt-2*. Predicted amino acid sequences are shown beneath the chromatograms.

(D) *Marchantia polymorpha* wild-type Tak-1 and Mp*npt-2* grown on MGRL medium for 16 days.

Scale bar: 1 cm.

(E) Quantification of thallus size in Tak-1 and Mp*npt-2*. $n$ = 4. $P$ values were determined by two-tailed Welch's $t$-test.

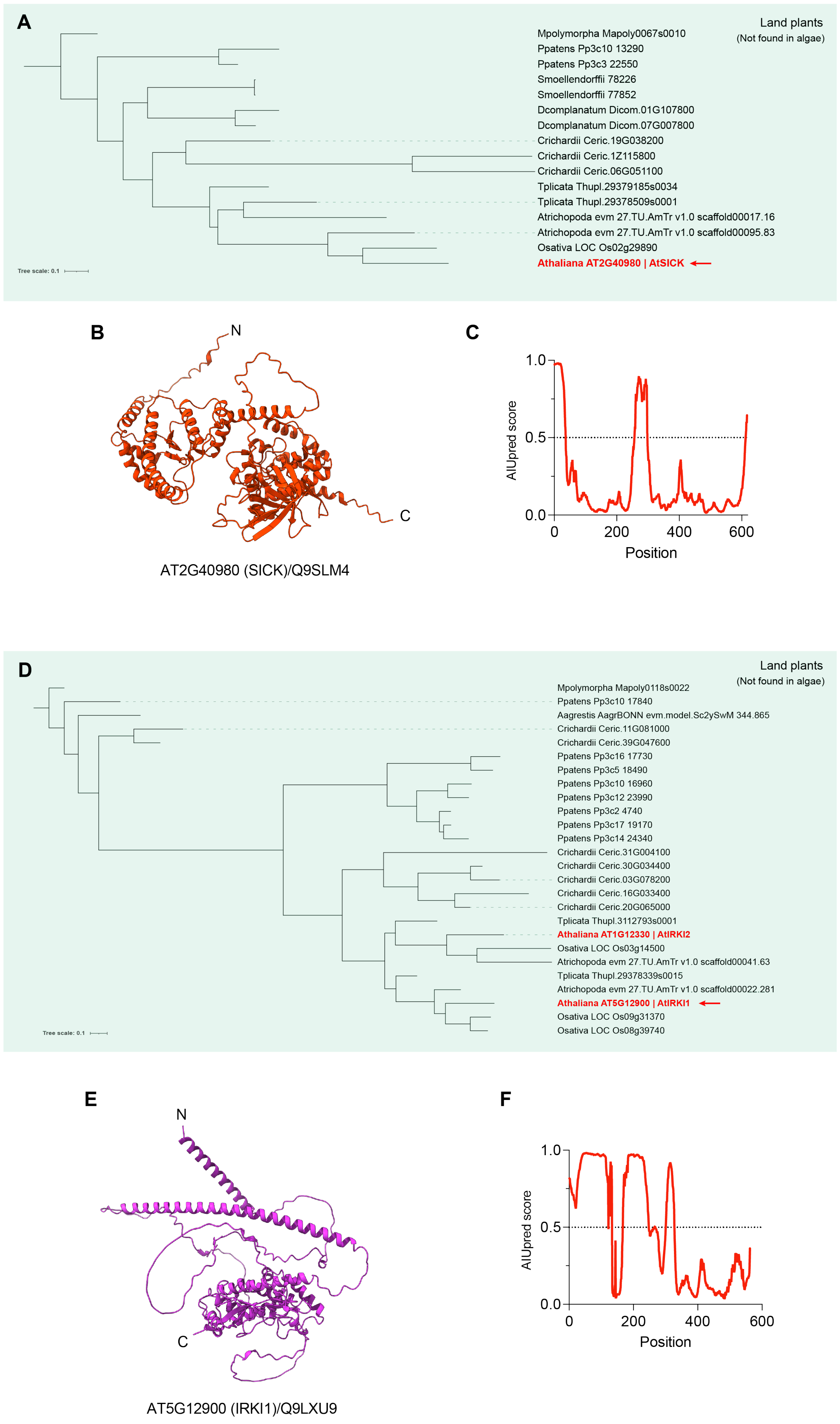

A
Land plants
(Not found in algae)
Mpolymorpha Mapoly0067s0010
Ppatens Pp3c10 13290
Ppatens Pp3c3 22550
Smoellendorffii 78226
Smoellendorffii 77852
Dcomplanatum Dicom.01G107800
Dcomplanatum Dicom.07G007800
Crichardii Ceric.19G038200
Crichardii Ceric.1Z115800
Crichardii Ceric.06G051100
Tplicata Thupl.29379185s0034
Tplicata Thupl.29378509s0001
Atrichopoda evm 27.TU.AmTr v1.0 scaffold00017.16
Atrichopoda evm 27.TU.AmTr v1.0 scaffold00095.83
Osativa LOC Os02g29890
Athaliana AT2G40980 | AtSICK
Tree scale: 0.1
B
N
C
AT2G40980 (SICK)/Q9SLM4
C
AIUpred score
1.0
0.5
0.0
0
200
400
600
Position
D
Land plants
(Not found in algae)
Mpolymorpha Mapoly0118s0022
Ppatens Pp3c10 17840
Aagrestis AagrBONN evm.model.Sc2ySwM 344.865
Crichardii Ceric.11G081000
Crichardii Ceric.39G047600
Ppatens Pp3c16 17730
Ppatens Pp3c5 18490
Ppatens Pp3c10 16960
Ppatens Pp3c12 23990
Ppatens Pp3c2 4740
Ppatens Pp3c17 19170
Ppatens Pp3c14 24340
Crichardii Ceric.31G004100
Crichardii Ceric.30G034400
Crichardii Ceric.03G078200
Crichardii Ceric.16G033400
Crichardii Ceric.20G065000
Tplicata Thupl.3112793s0001
Athaliana AT1G12330 | AtIRKI2
Osativa LOC Os03g14500
Atrichopoda evm 27.TU.AmTr v1.0 scaffold00041.63
Tplicata Thupl.29378339s0015
Atrichopoda evm 27.TU.AmTr v1.0 scaffold00022.281
Athaliana AT5G12900 | AtIRKI1
Osativa LOC Os09g31370
Osativa LOC Os08g39740
Tree scale: 0.1
E
N
C
AT5G12900 (IRKI1)/Q9LXU9
F
AIUpred score
1.0
0.5
0.0
0
200
400
600
Position

**Figure S17. Phylogenetic trees of SICK and IRKI1 homologs across eukaryotic lineages.**

(A) The phylogenetic tree was constructed using protein sequences homologous to the *Arabidopsis thaliana* SICK (AT2G40980) identified from representative green plant species. The query protein AtSICK is indicated by a red arrow. Protein sequences were aligned and analyzed using IQ-TREE3 with ultrafast bootstrap analysis (1,000 replicates). Bootstrap values are indicated at nodes. The tree was visualized using iTOL.

(B) Three-dimensional structure of SICK predicted by AlphaFold2.

(C) IDRs within SICK predicted by AIUPred.

(D) The phylogenetic tree was constructed using protein sequences homologous to the Arabidopsis thaliana IRKI1 (AT5G12900) identified from representative green plant species. The query protein AtIRKI1 is indicated by a red arrow. Protein sequences were aligned and analyzed using IQ-TREE3 with ultrafast bootstrap analysis (1,000 replicates). Bootstrap values are indicated at nodes. The tree was visualized using iTOL.

(E) Three-dimensional structure of IRKI1 predicted by AlphaFold2.

(F) IDRs within IRKI1 predicted by AIUPred.
